# Environmental context reveals a conditional role of the Tol-Pal system in envelope organization in *Acinetobacter baumannii*

**DOI:** 10.64898/2026.05.25.727595

**Authors:** Roberto Jhonatan Olea-Ozuna, Berenice Furlan, Suman Tiwari, Hanling Gong, Augusto C. Hunt-Serracin, Michael B. Whalen, Orietta Massida, Nicholas Dillon, Joseph M. Boll

## Abstract

Gram-negative bacteria must coordinate remodeling of the peptidoglycan cell wall with invagination of the outer membrane to preserve envelope integrity during growth and division. The conserved Tol-Pal system has been implicated in coordinating these processes, yet its physiological contribution to envelope organization remains unclear and may depend on environmental context. Here, we examined the role of Tol-Pal in coordinating envelope remodeling in *Acinetobacter baumannii* across distinct growth environments. Loss of Tol-Pal did not cause a major population growth defect, and septal peptidoglycan incorporation remained largely preserved under standard laboratory growth conditions. In contrast, under specific environmental conditions—including nutrient-rich media, altered osmotic conditions, and host-like environments—Tol-Pal deficiency disrupted the spatial organization of cell division and cell morphology. Tol-Pal mutants also exhibited modest but reproducible reductions in outer membrane barrier robustness and decreased fitness in environmental and host-associated contexts. Together, these findings demonstrate that Tol-Pal is not an essential component of the core division machinery but instead contributes to the coordinated organization of the Gram-negative envelope under conditions that impose additional physiological demands. More broadly, our results highlight how environmental context can reveal conditional roles for conserved envelope systems that are not apparent during standard laboratory growth.

**Importance:** The Gram-negative envelope is a complex, multilayered structure that must remain intact as cells grow and divide across diverse and often challenging environments. Coordination between peptidoglycan remodeling and outer membrane invagination is therefore critical for maintaining envelope organization and cellular fitness. Here, we show that the conserved Tol-Pal system in *Acinetobacter baumannii* contributes to the spatial organization of cell division and outer membrane robustness under specific environmental conditions. Although Tol-Pal deficiency permits sustained population growth under standard laboratory conditions, its absence disrupts envelope organization and compromises bacterial fitness in environmental and host-associated contexts. These findings demonstrate how environmental conditions can expose conditional roles for conserved envelope systems and highlight the importance of physiological context in shaping bacterial cell envelope organization.

## Introduction

The Gram-negative bacterial envelope is a multilayered structure whose integrity depends on coordinated interactions between the inner membrane (IM), the peptidoglycan (PG) cell wall, and the outer membrane (OM) (1–3). During cell growth and division, these layers must remodel in a synchronized manner to preserve envelope continuity as cells elongate, constrict, and separate (4, 5). Disruption of this coordination can compromise barrier function, alter cell morphology, and ultimately reduce bacterial fitness (6–8).

Although the envelope is often considered primarily as a permeability barrier due to the protective properties of the OM, it also functions as a dynamic structure that undergoes extensive remodeling during the bacterial cell cycle (9, 10). Septal PG synthesis and remodeling must occur in concert with invagination of the OM to maintain structural integrity during cytokinesis (11, 12). Multiple molecular systems have therefore been identified that link PG remodeling to OM dynamics during division (4, 13–15). Importantly, the contribution of these systems can depend on the physiological conditions in which cells grow, as envelope-associated defects often become evident when environmental conditions place additional mechanical or organizational demands on the cell surface (16–18).

One conserved system implicated in coordinating OM behavior with cell wall remodeling is the Tol-Pal complex (19–21). Tol-Pal forms a trans-envelope assembly composed of an IM motor complex formed by TolQ, TolR, and TolA that harnesses the proton motive force, together with the periplasmic protein TolB and the PG-binding OM lipoprotein Pal (22, 23), as illustrated in **Figure S1A**. In *Escherichia coli* and related Enterobacteria, Tol-Pal localizes to the division septum and contributes to OM invagination during cytokinesis (12, 20, 24). Loss of Tol-Pal leads to increased OM permeability, enhanced vesiculation, and division-associated morphological defects, consistent with a role in supporting envelope organization during growth (25, 26). In addition, recent studies have suggested that Tol-Pal contributes to maintenance of OM lipid homeostasis, indicating that this complex may play a broader role in stabilizing envelope architecture (27–29).

Despite these functions, Tol-Pal–associated phenotypes are often relatively modest under routine laboratory growth conditions (20, 24, 30, 31). This observation raises the possibility that the physiological contribution of Tol-Pal becomes more apparent in specific environmental contexts that impose additional demands on envelope organization. Importantly, the extent to which envelope coordination systems such as Tol-Pal contribute to cell division and envelope integrity across different physiological environments remains poorly defined.

*Acinetobacter baumannii* provides a tractable system to examine how environmental conditions shape envelope organization. *A. baumannii* persists across diverse environmental and host-associated settings, including hospital surfaces and infected tissues, where cells encounter osmotic fluctuations, desiccation, nutrient variation, and immune-associated stresses (32–34). Such environments likely impose variable and dynamic constraints on envelope stability during growth and division.

Here, we investigated how the conserved Tol-Pal system contributes to envelope coordination in *A. baumannii* across growth environments that differ in osmotic and physiological conditions. Using strains ATCC 17978 and the clinical isolate AB5075, we examined whether Tol-Pal–associated phenotypes are conserved across divergent genetic backgrounds. Our results show that Tol-Pal deficiency permits sustained population growth under standard laboratory conditions but disrupts the spatial organization of cell division in specific growth environments. In addition, loss of Tol-Pal modestly compromises OM robustness and reduces bacterial fitness under environmental and host-associated conditions. Together, these findings indicate that the contribution of Tol-Pal to envelope organization is strongly influenced by environmental context and becomes particularly important under conditions that impose additional demands on envelope coordination.

## Results

### Tol-Pal deficiency permits sustained population growth under standard laboratory conditions

To determine whether the Tol-Pal system contributes to basal growth in *A. baumannii*, we constructed in-frame deletion mutants lacking the entire *tol*–*pal* locus as well as mutants targeting the individual transcriptional units *tolQRA* and *tolB*–*pal* in the ATCC 17978 background (**Figure S1B**). These deletions disrupt the trans-envelope Tol-Pal complex by removing components spanning the IM, periplasm, and OM.

Population growth was monitored in commonly used laboratory media, including Luria-Bertani (LB), Tryptic Soy Broth (TSB), and Brain Heart Infusion (BHI) (**Figure S2A**). Across all three media, Δ*tol*–*pal*, Δ*tolQRA*, and Δ*tolB*–*pal* strains exhibited broadly similar growth dynamics to the wild-type strain. Although minor differences in growth kinetics were observed in some conditions, mutant cultures reached comparable population densities throughout the growth cycle, indicating sustained population expansion under nutrient-replete conditions.

To determine whether this behavior was conserved across genetic backgrounds, transposon insertion mutants disrupting *tolQ*, *tolR*, *tolA*, *tolB*, or *pal* were examined in the clinical isolate AB5075 (**Figure S2B**). As observed in ATCC 17978, AB5075 mutants displayed broadly similar growth patterns to the wild-type strain in LB, TSB, or BHI.

Together, these results show that disruption of the Tol-Pal system does not cause a major population growth defect under standard laboratory conditions, and that basal growth of *A. baumannii* proceeds largely independently of Tol-Pal.

### Tol-Pal deficiency causes environment-dependent defects in cell division organization

Having established that Tol-Pal deficiency permits sustained population growth under standard laboratory conditions, we next examined whether loss of Tol-Pal affects cellular architecture across distinct growth environments (**Figure 1**). Wild-type and Tol-Pal–deficient strains were analyzed by phase-contrast microscopy and fluorescent D-amino acid labeling (HADA), which is incorporated into the PG, to assess the spatial organization of PG incorporation during division.

**Figure 1.**
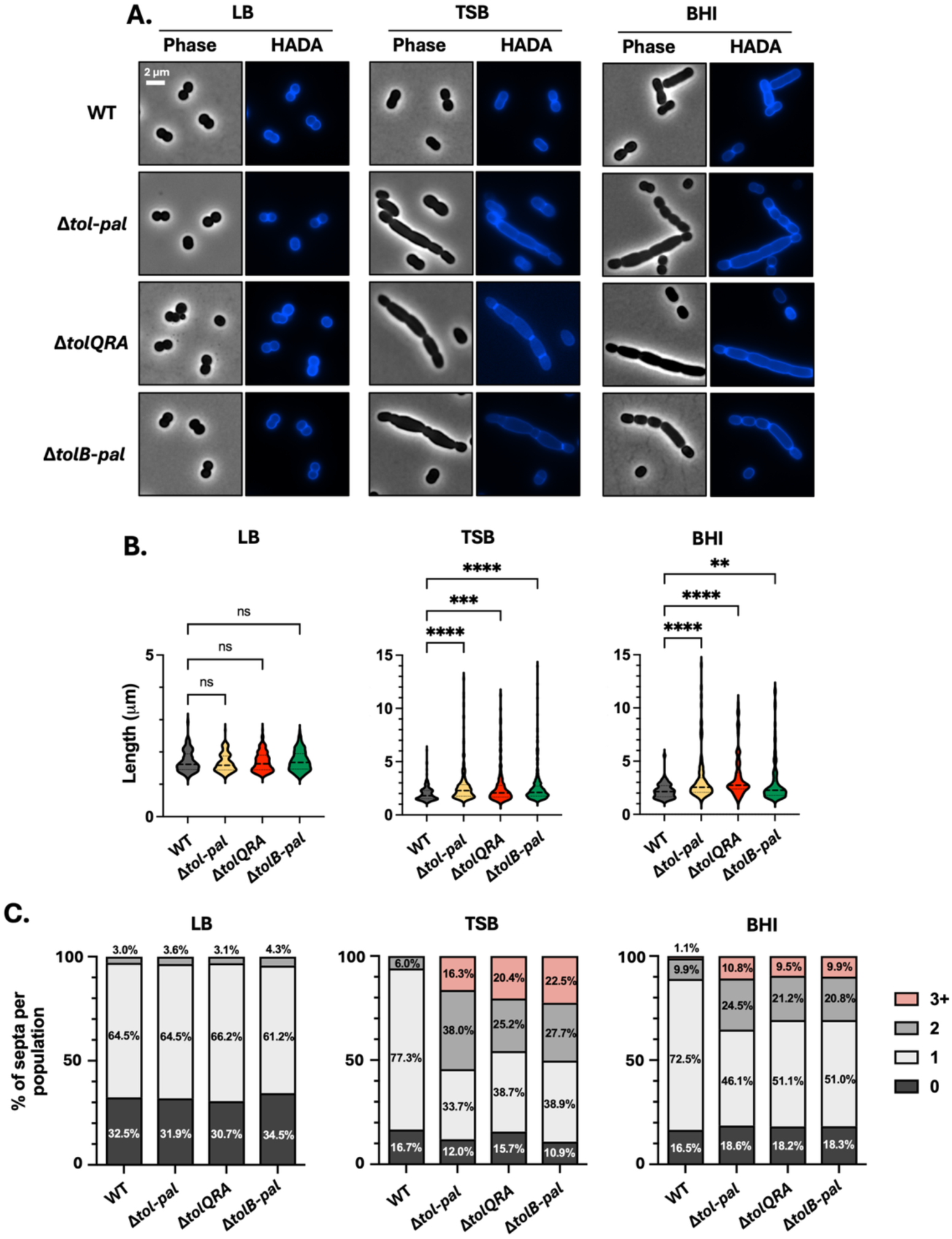
Tol-Pal deficiency causes environment-dependent defects in cell division organization. (A) Phase-contrast and fluorescent D-amino acid (HADA) labeling of wild-type (WT), Δ*tol–pal*, Δ*tolQRA*, and Δ*tolB–pal A. baumannii* ATCC 17978 cells grown in LB, TSB, or BHI to mid-exponential phase. Phase-contrast images show cell morphology, and HADA labeling marks sites of active PG synthesis. Scale bars, 2 µm. (B) Quantification of cell length distributions corresponding to panel A. For each condition, ∼250–300 cells were analyzed across independent fields of view across biological replicates. Statistical significance was assessed using ordinary one-way ANOVA. Error bars represent the standard deviation from three independent biological replicates. (C) Quantification of septa per cell based on HADA labeling patterns. Cells grown in TSB and BHI display increased filamentation, multiple septa, and mislocalized septal PG synthesis in Tol-Pal–deficient strains but not during growth in LB. For each condition, ∼250–300 cells were analyzed across independent fields of view.

During growth in LB, Tol-Pal mutants maintained a short rod-shaped morphology comparable to that of the wild-type strain (**Figure 1A**). Quantitative analysis of cell length distributions revealed no significant difference between wild-type and mutant populations under these conditions (**Figure 1B**). Consistent with this morphology, HADA labeling showed discrete mid-cell incorporation, indicating properly localized septal PG synthesis. Quantification of septa per cell further revealed similar distributions between wild-type and Tol-Pal–deficient populations when grown in LB media (**Figure 1C**).

In contrast, growth in TSB or BHI resulted in pronounced division-associated defects in Tol-Pal–deficient cells (**Figure 1A**). Cells frequently exhibited extensive filamentation and incomplete constrictions. Quantitative analysis revealed a significant increase in mean cell length and an increase in the proportion of elongated cells relative to wild-type **(Figure 1B**). Septation patterns were altered, with Tol-Pal mutants displaying a higher frequency of cells containing multiple septa compared with wild-type populations (**Figure 1C**).

Consistent with these observations, HADA labeling in Tol-Pal–deficient cells was frequently detected across multiple septal sites rather than focused at discrete mid-cell constrictions, and the fraction of cells displaying focused mid-cell incorporation was markedly reduced. These observations indicate that Tol-Pal contributes to the coordinated spatial organization of septum formation and OM invagination during division. Genetic complementation of the *tol*–*pal* mutants restored cell length distributions to near-wild-type levels in TSB and BHI media, confirming that the phenotypes are specifically attributable to loss of Tol-Pal (**Figure S3A**).

Comparable morphological defects were observed in the AB5075 background under the same growth conditions (TSB and BHI) (**Figure S3B**), where transposon mutants disrupting *tolQ*, *tolR*, *tolA*, *tolB*, or *pal* displayed increased cell length and irregular morphologies, indicating conservation of this phenotype across *A. baumannii* lineages.

To determine whether the division defects observed in rich media reflected specific nutrient composition or broader environmental properties, TSB and BHI were diluted 1:5 with water. Under these diluted conditions, filamentation in Δ*tol*–*pal* cells was substantially reduced (**Figure S3C**), with cell length distributions approaching those of the wild-type strain and restoration of focused septal HADA labeling. These observations suggest that the morphological defects observed in concentrated rich media are influenced by environmental conditions rather than strictly dependent on nutrient identity.

Together, these results show that Tol-Pal contributes to maintaining the spatial organization division-associated PG incorporation in a context-dependent manner.

### Osmotic conditions influence envelope organization in Tol-Pal mutants

The division defects observed in concentrated growth media and their suppression upon medium dilution suggested that physicochemical properties of the growth environment influence envelope organization in the absence of Tol-Pal. To examine whether osmotic conditions contribute to this phenotype, we examined cell morphology and growth under defined hypo-and hyperosmotic environments.

Standard LB Miller medium contains approximately 171 mM NaCl and therefore represents a condition of moderate osmolarity. To reduce osmolarity, strains were grown in LB lacking added NaCl (LB0N). Under these reduced-osmolarity conditions, Δ*tol*–*pal*, Δ*tolQRA*, and Δ*tolB*–*pal* strains exhibited pronounced filamentation and disrupted septal organization relative to wild-type (**Figure 2A**). Phase-contrast microscopy revealed elongated cells with irregular constrictions, and HADA labeling showed PG incorporation that was no longer spatially restricted to a discrete mid-cell septal plane. Genetic complementation restored normal cell morphology (**Figure S4A**).

**Figure 2.**
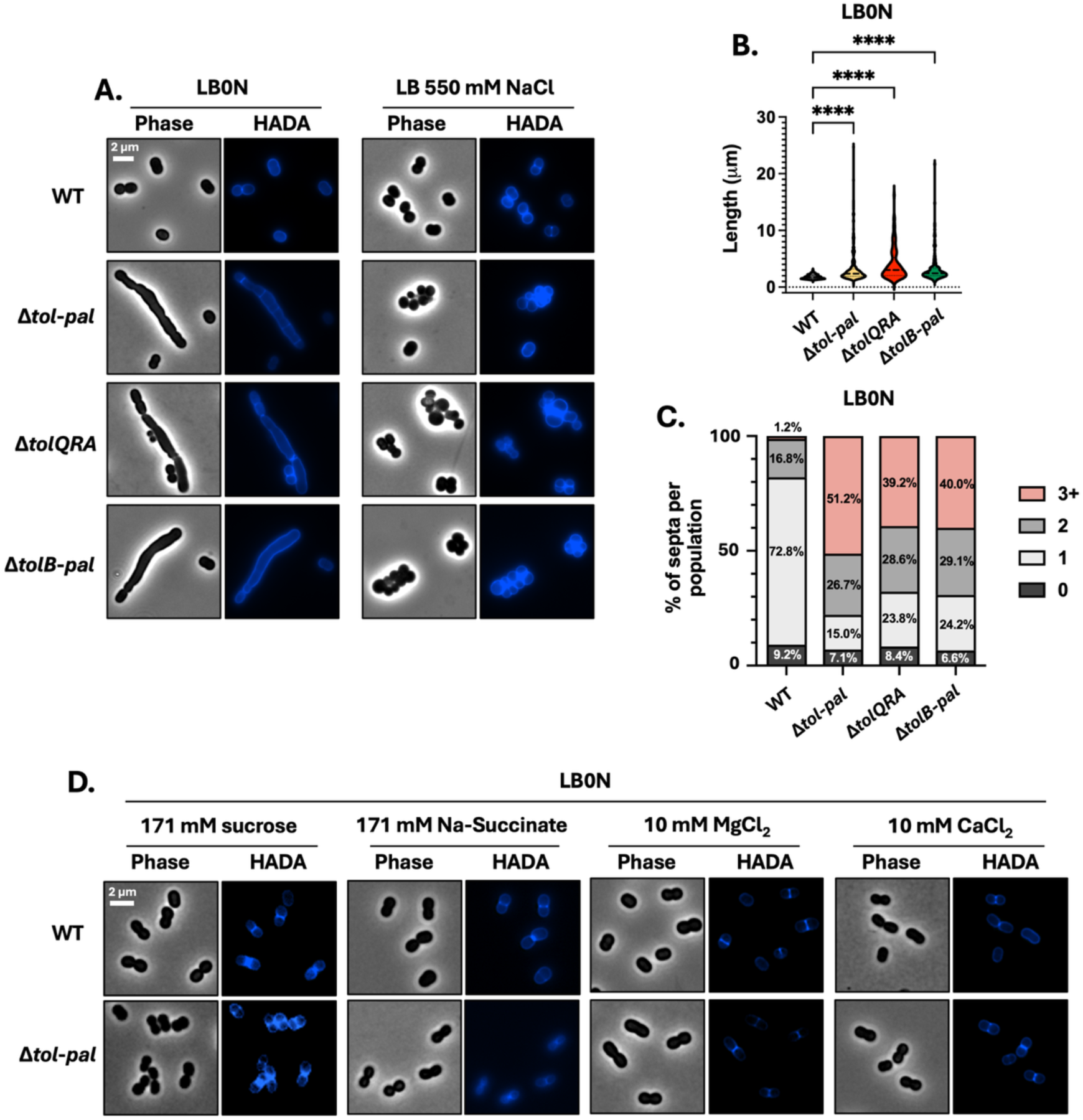
Osmotic stress disrupts envelope organization in Tol-Pal–deficient cells. Osmotic conditions influence envelope organization in the absence of Tol-Pal. (A) Phase-contrast and HADA labeling of WT, Δ*tol*–*pal*, Δ*tolQRA*, and Δ*tolB*–*pal* strains grown in LB without NaCl (LB0N). Tol-Pal–deficient cells exhibit filamentation and loss of focused mid-cell PG incorporation. In contrast, growth in LB supplemented with 550 mM NaCl induces cellular aggregation and aberrant PG organization in Tol-Pal–deficient cells. (B, C) Quantification of cell length distributions and number of septa per cell under hypo-and hyperosmotic conditions. For each condition, ∼250–300 cells were analyzed across independent fields of view. Statistical significance was assessed using ordinary one-way ANOVA. (D) Phase-contrast and HADA labeling of WT and Δ*tol*–*pal* cells grown in LB0N supplemented with osmoprotective solutes (sucrose or sodium succinate) or divalent cations (MgCl₂ or CaCl₂). Supplementation restores cell morphology and septal organization in Tol-Pal–deficient cells. Scale bars, 2 µm.

Quantitative analysis of cell length distributions confirmed a marked increase in elongated cells and cells containing multiple septa in Tol-Pal–deficient populations grown in LB0N relative to wild-type cells (**Figure 2B and 2C**).

Despite these morphological defects, Tol-Pal mutants maintained population growth in LB0N and reached optical densities approaching those of the wild-type strain at later time points (**Figure S4B**). These observations indicate that PG incorporation during division remains active in the absence of Tol-Pal, although its spatial organization becomes disrupted under reduced osmolarity.

To determine whether these phenotypes reflected sensitivity to osmotic imbalance, LB0N was supplemented with osmoprotective solutes or divalent cations. Addition of sucrose or sodium succinate, as well as MgCl₂ or CaCl₂, suppressed filamentation and restored focused septal PG labeling in Tol-Pal–deficient cells (**Figure 2D**).

We next examined hyperosmotic conditions by increasing NaCl to 550mM. Under these conditions, Tol-Pal mutants formed dense cellular aggregates and exhibited irregular patterns of PG incorporation (**Figure 2A**). Consistent with these morphological changes, mutant cultures also displayed delayed growth relative to the wild-type strain (**Figure S4C**). Genetic complementation restored normal cellular dispersion (**Figure S4A**).

Together, these results show that osmotic conditions influence envelope organization in Tol-Pal-deficient cells. In the absence of Tol-Pal, septal PG synthesis persists but coordinated envelope remodeling becomes disrupted under both hypo-and hyperosmotic conditions.

### Tol-Pal contributes OM barrier robustness

The division defects and osmotic sensitivity observed in Tol-Pal–deficient cells prompted us to examine whether loss of this system also affects OM barrier function. Susceptibility to detergents and antibiotics whose activity depends on OM permeability was evaluated in the ATCC 17978 background.

Under standard LB growth conditions, Tol-Pal mutants displayed increases in sensitivity to bile salts, deoxycholate, and the detergent–chelator combination SDS–EDTA relative to the WT strain (**Figure 3A**). Susceptibility to antibiotics with limited penetration across an intact OM, including vancomycin, bacitracin, and rifampicin, was similarly increased (**Figure 3B**). Although modest in magnitude, these differences were consistently observed across independent experiments, indicating reduced OM barrier robustness. Increased sensitivity to cationic antimicrobial peptides was also observed (**Figure 3B**) and was restored by genetic complementation (**Figure S5A**), confirming that these phenotypes result from loss of Tol-Pal.

**Figure 3.**
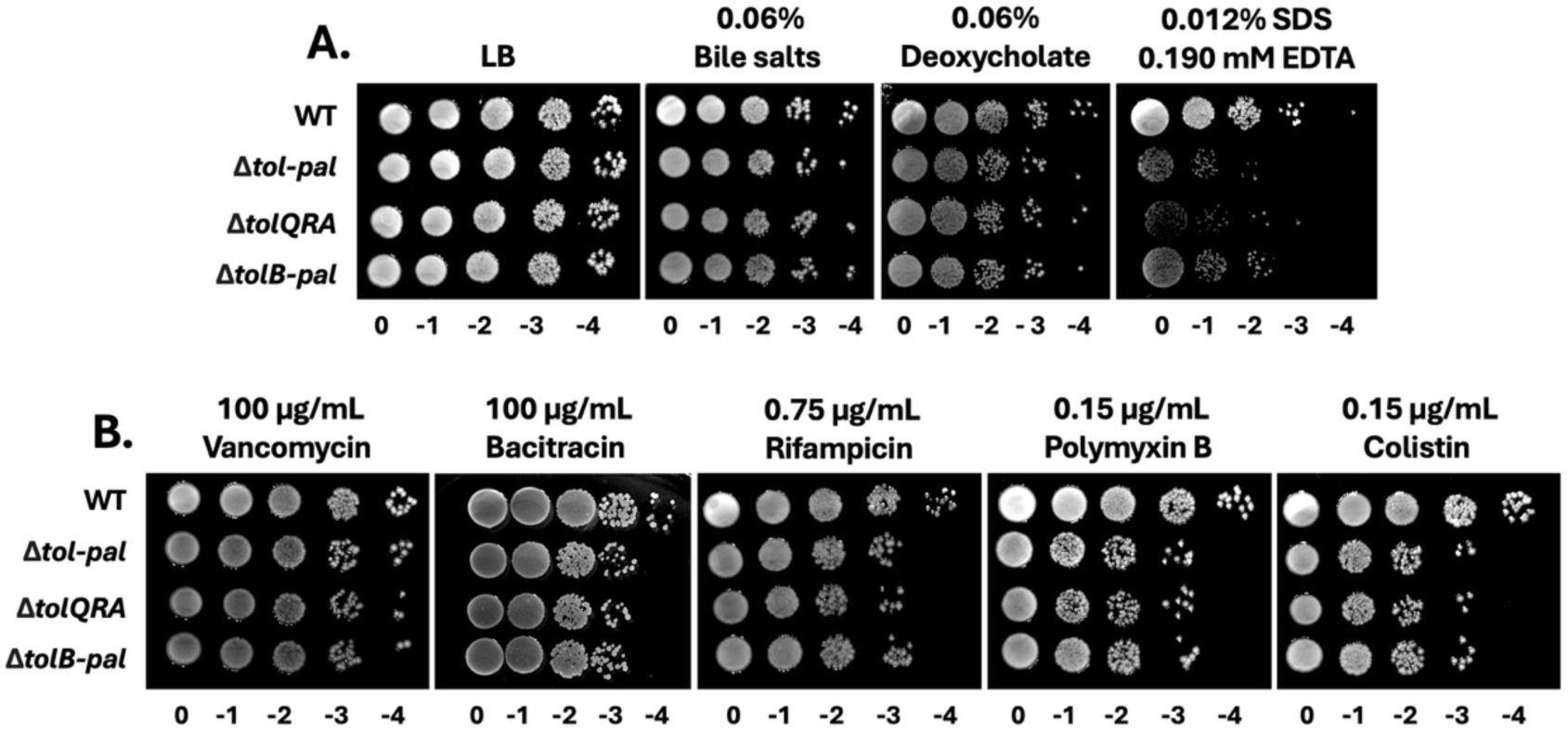
Tol-Pal contributes to OM barrier robustness. Loss of Tol-Pal compromises OM barrier function. (A) Spot-dilution assays of WT, Δ*tol–pal*, Δ*tolQRA*, and Δ*tolB–pal* strains grown on LB agar containing bile salts, deoxycholate, or SDS–EDTA. (B) Spot-dilution assays assessing susceptibility to antibiotics with limited penetration across an intact OM, including vancomycin, bacitracin, rifampicin, polymyxin B, and colistin. Tol-Pal–deficient strains show increased sensitivity relative to WT. All assays were performed under standard LB conditions.

Comparable increases in sensitivity to SDS–EDTA and colistin were observed in AB5075 transposon mutants disrupting individual Tol-Pal components (**Figure S5B**), indicating that reduced OM robustness is conserved across *A. baumannii* lineages.

To determine whether these barrier defects were associated with alterations in envelope lipid composition, we examined phospholipid and lipooligosaccharide (LOS) profiles in Tol-Pal mutants (**Figure S6**). Phospholipids were labeled with ³²P during growth, extracted, and analyzed by thin-layer chromatography (TLC). The major phospholipid species, including phosphatidylethanolamine (PE), phosphatidylglycerol (PGL), and cardiolipin (CL), were present at comparable levels in wild-type and Tol-Pal–deficient strains in both ATCC 17978 and AB5075 backgrounds (**Figure S6A**). Phospholipids were quantified in **Table S1**. Similarly, analysis of LOS by SDS-PAGE revealed similar banding patterns between wild-type and Tol-Pal mutants (**Fig. S6B**). LOS was quantified in **Table S2**.

These results indicate that loss of Tol-Pal does not substantially alter global phospholipid composition or LOS profiles, suggesting that the barrier defects observed here arise from impaired envelope organization rather than changes in lipid composition.

Together, these findings indicate that Tol-Pal contributes to maintaining OM barrier robustness even though overall envelope lipid composition remains largely unchanged.

### Tol-Pal supports bacterial fitness and virulence in environmental and host-associated contexts

Because Tol-Pal contributes to envelope organization and OM robustness, we next examined whether its loss affected bacterial fitness under environmental and host-associated conditions.

In the ATCC 17978 background, Tol-Pal mutants exhibited decreased survival during prolonged desiccation (**Figure 4A**), reduced biofilm formation (**Figure 4B**), and impaired surface-associated motility (**Figure 4C**). Complementation of the *tol*-*pal* locus restored biofilm formation and motility to near-wild-type levels (**Figure S7A** and **S7B**). Reduced motility was likewise observed in AB5075 transposon mutants (**Figure S7C**), indicating conservation of this phenotype across genetic backgrounds.

**Figure 4.**
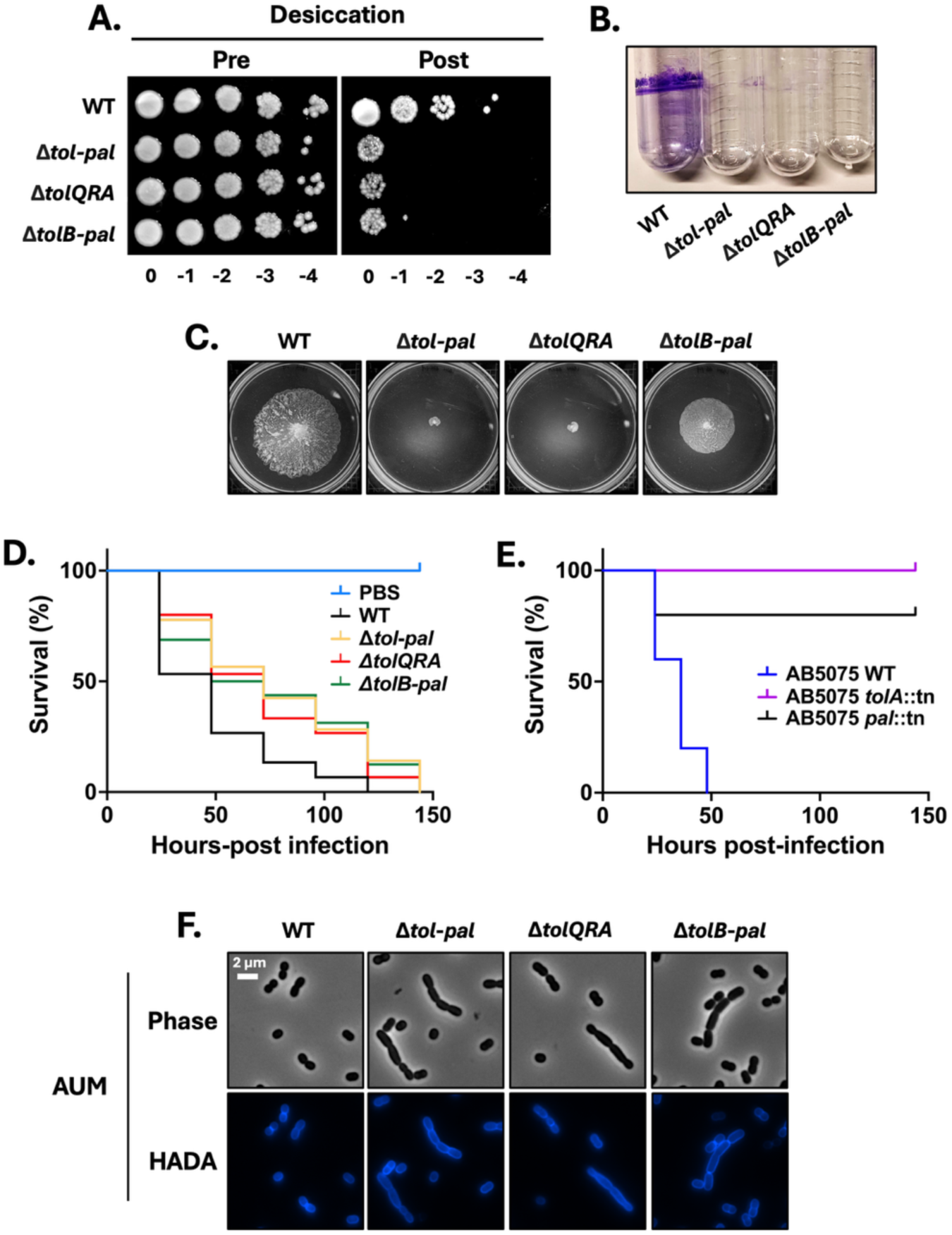
Tol-Pal supports bacterial fitness in environmental and host-associated contexts. Tol-Pal supports envelope-dependent fitness traits in environmental and host-associated conditions. (A) Survival of WT and Tol-Pal–deficient strains following prolonged desiccation on abiotic surfaces. (B) Quantification of biofilm formation by WT and Tol-Pal–deficient strains. (C) Surface-associated motility assessed on LB0N soft-agar plates. (D) Survival of *Galleria mellonella* larvae infected with WT or Δ*tol–pal* strains (*n* = 25 larvae per group). (E) Survival of mice inoculated with WT or AB5075 Δ*tolA* or Δ*pal* transposon mutants in a murine pneumonia model (*n* = 5 animals per group). (F) Phase-contrast and HADA labeling of WT and Tol-Pal–deficient strains grown in artificial urine medium (AUM), showing altered cell morphology and septal organization despite sustained population growth. Scale bars, 2 µm.

We next examined host-associated fitness. Tol-Pal mutants displayed reduced virulence in a *Galleria mellonella* infection model (**Figure 4D**), as reflected by increased host survival relative to infections with the wild-type strain. Consistently, disruption of *tolA* or *pal* in the AB5075 background reduced virulence in a murine pneumonia model (**Figure 4E**).

To further examine Tol-Pal function in a physiologically relevant environment, we assessed cell morphology and growth in a medium that mimics the composition of human urine (Artificial Urine Medium; AUM). Under these conditions, Tol-Pal mutants in both the ATCC 17978 (Δ*tol*-*pal*, Δ*tolQRA*, and Δ*tolB*-*pal*) (**Figure 4F**) and AB5075 (*tolA* and *pal* transposon) (**Figure S7D**) backgrounds exhibited altered cell morphology and defects in septal organization relative to their respective wild-type strains. Despite these morphological abnormalities, Tol-Pal mutants displayed population growth comparable to that of the wild-type strain (**Figure S7E**). These observations indicate that Tol-Pal contributes to maintaining coordinated envelope architecture in host-like environments despite sustained population growth.

Collectively, these findings suggest that although Tol-Pal deficiency permits sustained population growth under standard laboratory conditions, it compromises several envelope-dependent traits that contribute to bacterial fitness in environmental and host-associated contexts. Together, these results show that Tol-Pal supports envelope stability and physiological adaptability in *A. baumannii*.

## Discussion

The Gram-negative cell envelope is an integrated structure whose integrity depends on coordinated remodeling of the PG cell wall and OM during cell growth and division (1, 35). The Tol-Pal system has been extensively characterized in *E. coli* and related bacteria, where it contributes to OM invagination during cytokinesis, maintenance of OM integrity, and resistance to envelope stress (20, 36). Although Tol-Pal participates in these processes, its physiological contribution can vary across organisms and growth environments.

In *A. baumannii*, previous studies have linked Tol-Pal to cell morphology, antibiotic susceptibility, and virulence, suggesting that this complex participates in envelope-associated processes (37). Our findings refine this view by defining the environmental contexts in which Tol-Pal contributes to coordinated envelope remodeling, OM robustness, and bacterial fitness.

We show that Tol-Pal deficiency permits sustained population growth and preserves division-associated PG incorporation across multiple laboratory media. However, in specific growth environments, including nutrient-rich media such as TSB and BHI, conditions that alter osmotic balance, and host-like environments such as AUM, the spatial organization of septation and normal cell morphology become dependent on Tol-Pal. Septal PG synthesis persists in its absence, as indicated by continued incorporation of fluorescent D-amino acids, yet its spatial organization becomes disrupted. These observations indicate that Tol-Pal is not required for initiation of division itself, but instead contributes to coordination between envelope layers during cytokinesis under specific environmental conditions (12).

The contrasting responses of Tol-Pal–deficient cells under hypo-and hyperosmotic conditions further highlight the context-dependent role of Tol-Pal in envelope organization. Under reduced osmolarity, mutants exhibit filamentation and septal PG labeling that was no longer spatially restricted, which was suppressed by osmoprotectants and divalent cations. Under elevated osmolarity, growth delay and cellular aggregation become evident. Together, these phenotypes indicate that Tol-Pal contributes to envelope organization in a context-dependent manner rather than functioning as an obligate structural component of the core division machinery. Similar conditional phenotypes have been reported in Enterobacteria, suggesting that environmental sensitivity of envelope coordination systems may represent a conserved physiological principle (27, 30). In contrast to observations in *E. coli*, where Tol-Pal mutants exhibit temperature-sensitive growth and division defects at elevated temperatures (12), Tol-Pal-deficient *A. baumannii* strains did not display major growth or morphological defects at 42 °C under the conditions tested (data not shown). These observations suggest that while environmental sensitivity may represent a conserved feature of Tol-Pal disruption, the specific environmental conditions that expose these defects can differ across Gram-negative species.

Beyond division-associated defects, Tol-Pal deficiency produces modest but reproducible reductions in OM barrier robustness. Increased susceptibility to detergents, bile salts, and antibiotics with limited OM permeability indicates that envelope integrity is measurably altered even when population growth is maintained (19, 21). Although subtle, these defects likely lower the threshold at which additional environmental or host-imposed stresses compromise cellular performance.

Recent work has linked the Tol-Pal system to maintenance of OM lipid homeostasis in Gram-negative bacteria (27–29). Altered phospholipid distribution and increased vesiculation have been reported in Tol-Pal–deficient cells, suggesting that this complex contributes to stabilization of OM architecture. In the present study, global phospholipid and LOS profiles remained largely unchanged in Tol-Pal mutants, indicating that the barrier defects observed here are unlikely to arise from major alterations in lipid composition. Because these measurements reflect total cellular phospholipids, they do not resolve potential changes in lipid partitioning between the inner and outer membranes (38). Future studies examining membrane-resolved lipid composition will therefore be important to further define the contribution of Tol-Pal to envelope lipid organization. Even modest disruption of envelope architecture may nevertheless influence bacterial performance under physiologically relevant stresses.

Consistent with this interpretation, loss of Tol-Pal reduces fitness under conditions relevant to environmental persistence and infection, including survival to desiccation, surface-associated behaviors, and virulence in infection models. These findings reinforce the principle that envelope coordination systems may appear dispensable during routine laboratory growth yet become important determinants of bacterial fitness when cells encounter environmental or host-associated challenges.

Collectively, our results establish Tol-Pal as a conditionally important contributor to envelope coordination during cell division in *A. baumannii*. Rather than acting as an essential component of the division machinery, Tol-Pal contributes to maintaining spatial organization of envelope remodeling under nutrient-rich, osmotic, and host-like conditions. More broadly, these findings illustrate how physiological environments can reveal functional roles of conserved envelope systems that remain cryptic during routine laboratory growth, highlighting the importance of environmental context in shaping bacterial envelope organization and fitness.

## Acknowledgments

We thank members of the De Nisco laboratory for providing and advising on the use of artificial urine medium (AUM). The work was supported by funding from the National Institutes of Health (grants R35GM143053 to J.M.B) and (grant1R21AI190814-01A1 to N.D.). The funders had no role in study design, data collection and analysis, decision to publish, or preparation of the manuscript.

## Materials and Methods

### Bacterial strains and growth conditions

All bacterial strains and plasmids used in this study are listed in Table S2, and primer sequences are provided in Table S3. *A. baumannii* ATCC 17978 was used as the primary strain background unless otherwise indicated. Where specified, experiments were also performed in *A. baumannii* AB5075 to assess the generality of Tol-Pal–associated phenotypes across divergent lineages.

Strains were recovered from −80 °C glycerol stocks on Luria–Bertani (LB Miller) agar and incubated at 37 °C. Unless otherwise specified, *A. baumannii* was cultured aerobically at 37 °C in LB Miller broth (containing 171 mM NaCl), LB without added NaCl (LB0N), Tryptic Soy Broth (TSB), or Brain Heart Infusion (BHI). Liquid cultures were grown with shaking aeration (200 rpm).

Where indicated, LB0N was supplemented with sucrose (171 mM), sodium succinate (171 mM), NaCl (550 mM), MgCl₂ (10 mM), or CaCl₂ (10 mM). Antibiotics were used at the following final concentrations: kanamycin (15 or 30 µg mL⁻¹) and tetracycline (5 or 10 µg mL⁻¹).

*A. baumannii* strain ATCC 17978 has been shown to be a mixture of two variants (39). Colony PCR using primers specific to the cardiolipin synthase gene (*clsC2*) resulted in a PCR product, indicating this study was specific to only the *A. baumannii* 17978 UN variant. *A. baumannii* strains AB5075 has been shown to produce colonies with two opacity phenotypes that are phase variable (40). Microscopic examination showed most opaque forms in wild-type AB5075 with only 2-3 translucent colonies out of > 200 colonies found on each plate (*n* = 3 for each strain).

### Artificial urine medium

Artificial urine medium (AUM) was used to approximate the ionic composition of the urinary tract environment. A base artificial urine medium (bAUM) was prepared in distilled water containing the following components per 350 mL: peptone L37 (1 g), yeast extract (100 mg), sodium bicarbonate (2.1 g), sodium chloride (5.2 g), sodium sulfate·10H₂O (3.2 g), potassium dihydrogen phosphate (0.95 g), dipotassium hydrogen phosphate (1.2 g), and ammonium chloride (1.3 g). The solution was mixed thoroughly and sterilized by filtration (0.22 µm).

A 2× AUM formulation was generated by supplementing sterile bAUM with L-lactic acid (100 mg L⁻¹), urea (2.5 g L⁻¹), citric acid (500 mg L⁻¹), creatinine (400 mg L⁻¹), FeSO₄·7H₂O (100 mg L⁻¹), uric acid (70 mg L⁻¹), CaCl₂·2H₂O (500 mg L⁻¹), and MgSO₄·7H₂O (500 mg L⁻¹). The pH was adjusted to 6.5 using 1 M HCl and the medium was filter sterilized.

Working 1× AUM was prepared immediately prior to use by diluting the 2× solution 1:1 with sterile distilled water.

### Construction of mutant and complemented *A. baumannii* strains

Chromosomal deletion mutants in *A. baumannii* ATCC 17978 were generated using established Rec_Ab_-mediated recombineering protocols (41–43). Briefly, strains carrying the recombinase plasmid pMMB67EH-TetR::Rec_Ab_ were grown in LB broth supplemented with tetracycline (10 µg mL⁻¹) to an OD₆₀₀ of ∼0.05. Expression of Rec_Ab_ was induced with 2 mM IPTG at 37 °C 45 min after inoculation. Cells were harvested at mid-exponential phase (OD₆₀₀ ≈ 0.4), washed five times with ice-cold 10% glycerol, and concentrated for electroporation.

Linear PCR products containing a kanamycin resistance cassette flanked by FRT sites and gene-specific homology arms were introduced by electroporation. Following a 4 h recovery in LB containing IPTG (2 mM), transformants were selected on LB agar supplemented with kanamycin (15 µg mL⁻¹). The recombinase plasmid was cured using 2 mM NiCl₂ as previously described (44). The kanamycin cassette was subsequently excised using a FLP recombinase–encoding plasmid (pMMB67EH-TetR::FLP) induced with 1 mM IPTG, and successful excision was confirmed by PCR.

For genetic complementation, genes were cloned into pMMB67EH-Kan and expressed following induction with 1 mM IPTG. AB5075 transposon insertion mutants were obtained from the Manoil library and maintained with tetracycline (5 µg mL⁻¹) (45).

### Growth kinetics and cell morphology

Overnight cultures grown at 37 °C were diluted to an initial OD_600_ of approximately 0.05 in fresh medium. For growth curve measurements, 100 µL aliquots were transferred to a 96-well microplate and incubated at 37 °C with continuous shaking. OD_600_ was recorded at hourly intervals for 12 hours using a microplate reader.

For morphological analysis, mid-exponential phase cultures (OD₆₀₀ = 0.4–0.5) were labeled with 100 µM HADA (7-hydroxycoumarin-3-carboxylic acid–D-alanine) for 15 min to visualize active PG synthesis (46). Cells were fixed with 4% paraformaldehyde, washed with PBS, and mounted on 1.5% agarose pads prepared in PBS. Images were acquired using a Nikon Eclipse Ti-2 wide-field epifluorescence microscope equipped with a 100×/1.45 NA objective and a Photometrics Prime 95B camera. Quantitative analysis of cell length and HADA signal was performed using ImageJ/Fiji with the MicrobeJ plugin.

### Detergent and antibiotic sensitivity assays

Envelope integrity was assessed using spot-dilution assays. Mid-exponential cultures were normalized to an OD₆₀₀ of 0.05, serially diluted (10⁰–10⁻⁴), and spotted onto LB agar supplemented with bile salts (0.06%), sodium deoxycholate (0.06%), or SDS/EDTA combinations (0.01% SDS/0.175 mM EDTA or 0.012% SDS/0.190 mM EDTA).

Antibiotic susceptibility was evaluated using vancomycin (100 µg mL⁻¹), bacitracin (100 µg mL⁻¹), rifampicin (0.75 µg mL⁻¹), polymyxin B (0.15 µg mL⁻¹) or colistin (0.15 µg mL⁻¹). Plates were incubated at 37 °C for 18 h prior to imaging.

### Desiccation assays

Desiccation tolerance was assessed on abiotic surfaces as previously described (47), with minor modifications. Exponential-phase cultures were normalized in LB broth to an OD₆₀₀ of 0.1. Ten-microliter aliquots (∼1 × 10⁶ CFU) were spotted onto sterile 96-well polystyrene plates. Control (pre-desiccation) samples were immediately resuspended in 90 µL LB, serially diluted, and plated to determine initial viable counts.

Experimental samples were maintained in a climate-controlled chamber at 25 °C and 40% relative humidity. After 72 h, desiccated spots were rehydrated by vigorous resuspension in 90 µL LB, and viable cells were quantified by serial dilution and plating.

### Surface motility and biofilm assays

Surface-associated motility was evaluated on LB0N soft-agar plates (0.3% w/v agar) (48, 49). A single isolated colony was inoculated into the center of each plate using a sterile toothpick. Plates were prepared 24 h in advance and equilibrated at room temperature to standardize moisture content, then incubated statically at 37 °C for 12 h. Motility zones were documented by digital imaging.

Biofilm formation was assessed using both qualitative and quantitative assays (48, 49) in LB0N. For visualization of biofilm rings, 2 mL aliquots of normalized cultures (OD₆₀₀ = 0.05) were incubated statically in 5 mL polystyrene tubes at 37 °C for 48 h. Quantitative biofilm assays were performed in parallel using 96-well microtiter plates, with 100 µL of normalized cultures incubated under identical conditions. In both assays, planktonic cells were removed, adherent biomass was washed three times with sterile distilled water, and biofilms were stained with 0.1% (w/v) crystal violet for 15 min. After washing, bound dye was solubilized in 30% (v/v) acetic acid and quantified by measuring absorbance at 550 nm.

### *In vivo* virulence models

The *G. mellonella* virulence assay was performed as previously described (50). *G. mellonella* larvae (Vanderhorst, Inc., St. Marys, OH) were stored in the dark and used within seven days of receipt. Twenty-five larvae (200–300 mg) were randomly assigned to each experimental group. Bacterial inoculums were prepared from overnight cultures of *A. baumannii* ATCC 17978 grown in LB broth, washed, and resuspended in sterile PBS. Each larva was injected with ∼1 × 10⁶ CFU into the last left proleg. PBS-injected larvae served as sham controls. Larvae were incubated at 37 °C and monitored daily for up to 6 days; death was scored as lack of response to gentle stimulation.

For the murine pneumonia model, female C57BL/6 mice (7–8 weeks old; *n* = 5 per group) were anesthetized with ketamine/xylazine (100/10 mg kg⁻¹) and intratracheally inoculated with 1 × 10⁸ CFU of *A. baumannii* AB5075 in 40 µL sterile PBS. Mice were monitored every 12 hours for survival for 6 days. All animal experiments were performed in accordance with protocols approved by the Institutional Animal Care and Use Committee (IACUC) at the University of Texas at Dallas.

### Analysis of ³²P-labeled phospholipids

PL composition was analyzed as described (51–53) with minor modifications. To compare phospholipid composition among strains, phospholipids were metabolically labeled with [³²P]-orthophosphate and analyzed by thin-layer chromatography (TLC). Overnight cultures were diluted to an OD_600_ of approximately 0.05 in 5 mL LB broth containing 5 µCi mL⁻¹ [³²P]-orthophosphate (PerkinElmer) and grown at 37 °C with aeration for 3 h. Cells were harvested by centrifugation, and total lipids were extracted using the Bligh and Dyer method.

Radiolabeled phospholipids were separated by TLC on silica gel 60 plates using a solvent system consisting of chloroform, methanol, and acetic acid (65:25:5, v/v). After chromatography, plates were dried and exposed to a phosphorimaging screen. Radiolabeled species were visualized using an Amersham Typhoon FLA 9500 phosphorimager (GE Healthcare). Individual phospholipid species were quantified using ImageJ software, and the abundance of each lipid was calculated as a percentage of the total radiolabeled phospholipid signal per lane.

### Lipooligosaccharide (LOS) analysis

LOS profiles were assessed as described (54) with modifications. LOS profiles were analyzed by SDS-PAGE followed by carbohydrate-specific staining. Overnight cultures were diluted to an OD_600_ of approximately 0.05 in LB broth and grown at 37 °C with aeration until reaching mid-exponential phase (OD_600_ ≈ 1.0). Cells were harvested by centrifugation, and pellets were resuspended in 1× LDS sample buffer containing β-mercaptoethanol. Samples were boiled for 10 min to solubilize cellular components.

Following cooling, proteinase K was added and samples were incubated at 55 °C overnight to digest cellular proteins. Samples were then boiled again for 5 min and separated by SDS-PAGE. Gels were subsequently fixed and stained using the Pro-Q Emerald 300 Lipopolysaccharide Gel Stain Kit (Thermo Fisher Scientific) according to the manufacturer’s instructions to visualize LOS species.

---

**Figure S1.**
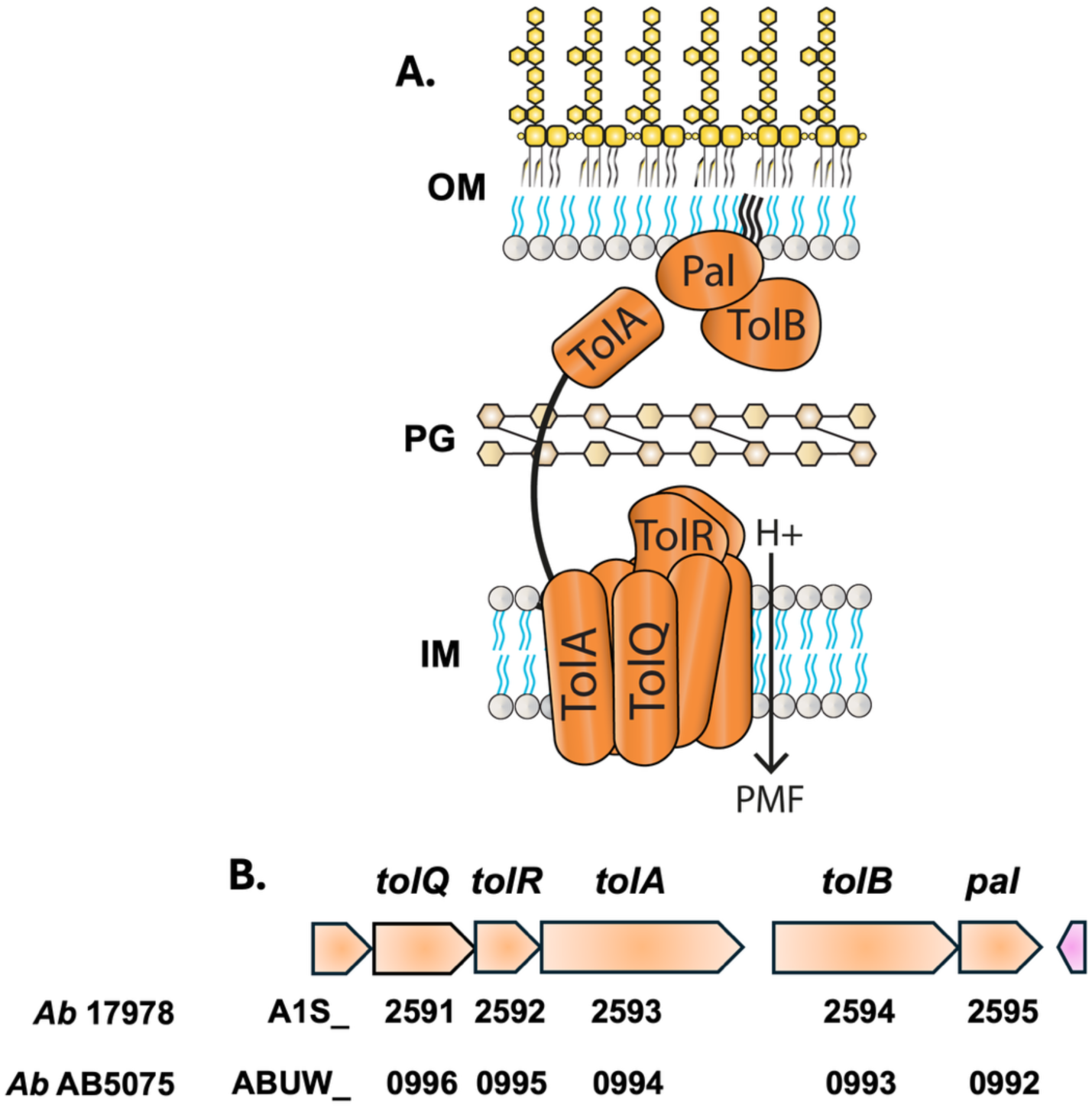
Organization of the Tol-Pal system in *A. baumannii*. (A) Schematic representation of the Tol-Pal trans-envelope complex showing the inner membrane (IM) motor components TolQ, TolR, and TolA, which harness the proton motive force (PMF), the periplasmic protein TolB, and the outer membrane (OM) lipoprotein Pal, which associates with peptidoglycan (PG). (B) The corresponding genetic organization of the *tol*–*pal* locus is, with gene annotations indicated below each locus. The locus is organized into two transcriptional units, *tolQRA* and *tolB–pal*, with gene annotations indicated below each locus.

**Figure S2.**
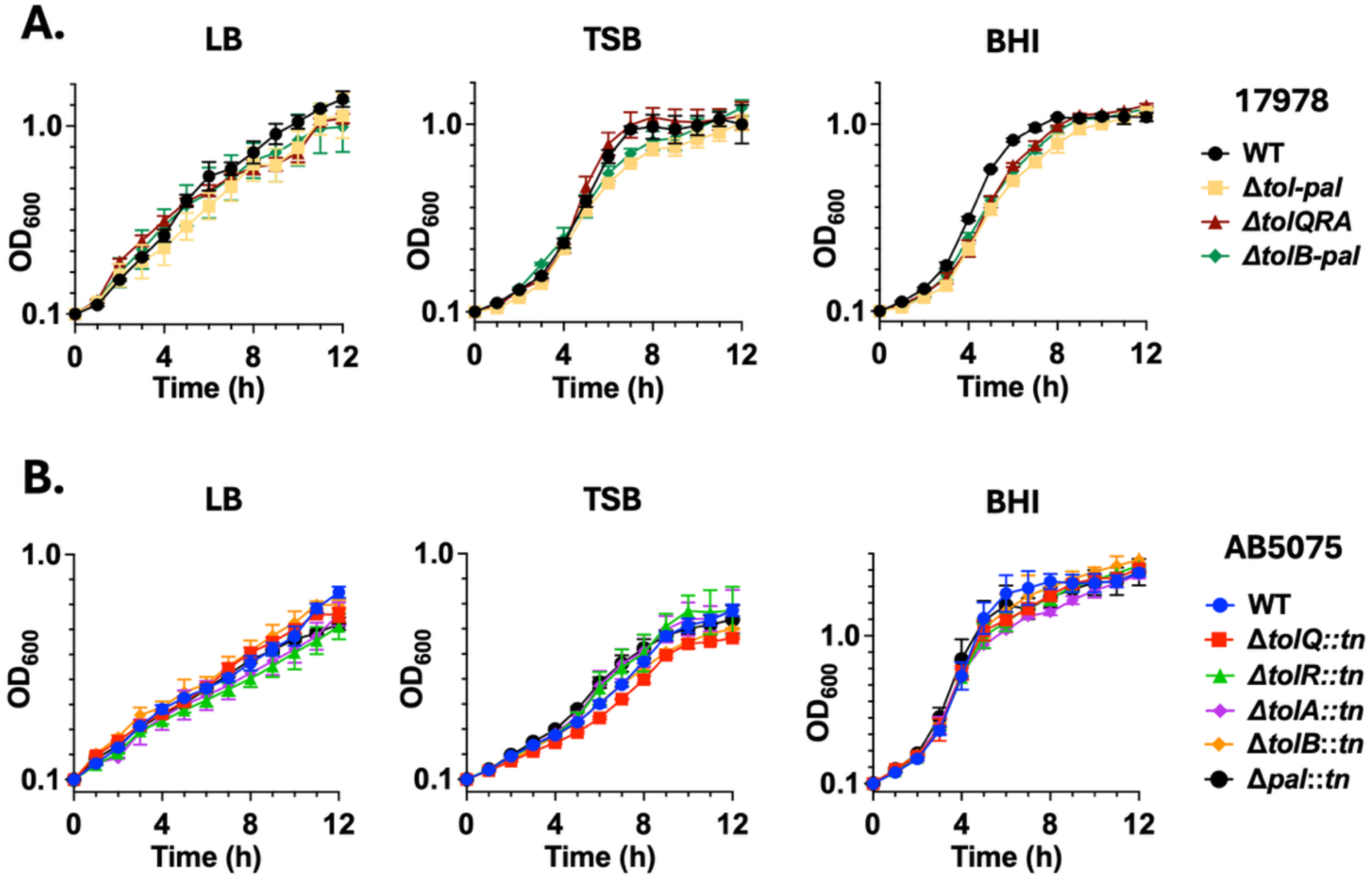
Tol-Pal deficiency permits sustained population growth under standard laboratory conditions. (A) Growth of wild-type (WT), Δ*tol–pal*, Δ*tolQRA*, and Δ*tolB–pal* generated in strain ATCC 17978 strains grown in LB, TSB, and BHI. (B) Growth of strain AB5075 WT and transposon mutants disrupting individual Tol-Pal components (*tolQ*, *tolR*, *tolA*, *tolB*, or *pal*) in the same media. Optical density was monitored over time. Error bars represent the standard deviation from three independent biological replicates.

**Figure S3.**
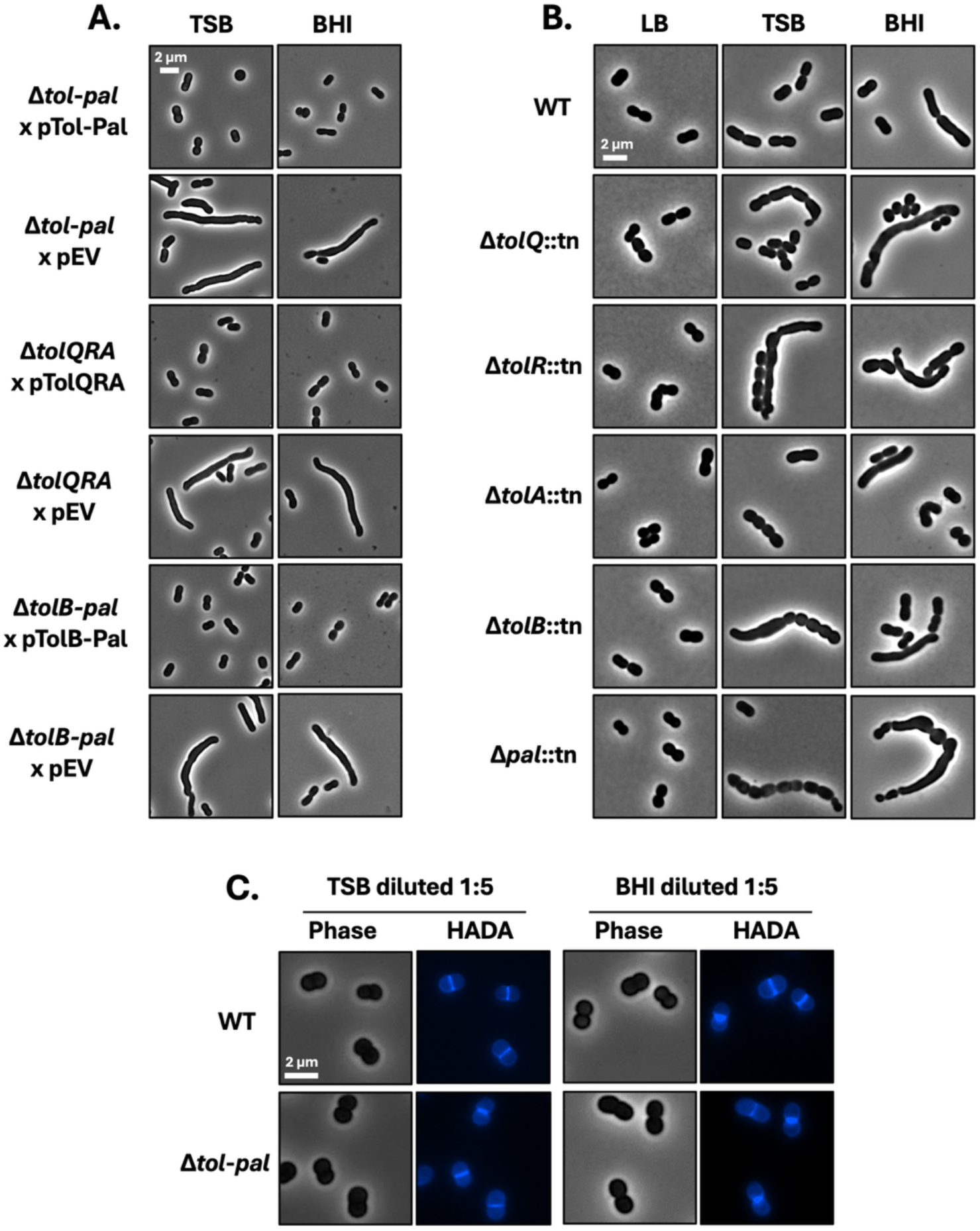
Genetic complementation restores normal cell morphology, and environmental modulation suppresses Tol-Pal–associated division defects. Role of Tol-Pal is conserved across genetic backgrounds. (A) Phase-contrast microscopy of *A. baumannii* ATCC 17978 *tol*-*pal* mutants and complemented strains grown in TSB or BHI. Cell length distributions are shown for complemented Δ*tol*–*pal*, Δ*tolQRA*, and Δ*tolB*–*pal* strains expressing the corresponding genes, compared with empty vector controls. Complementation restores wild-type cell morphology and reduces filamentation. (B) Phase-contrast microscopy of AB5075 Tol-Pal transposon mutants grown in LB, TSB, and BHI, showing conserved morphology defects across strain backgrounds and growth conditions. (C) Environmental modulation suppresses Tol-Pal–associated division defects. Phase-contrast and HADA labeling of WT and Δ*tol*–*pal* cells grown in TSB or BHI diluted 1:5 with water. Medium dilution suppresses filamentation and restores focused mid-cell PG incorporation in Δ*tol*–*pal* cells. Scale bars, 2 µm.

**Figure S4.**
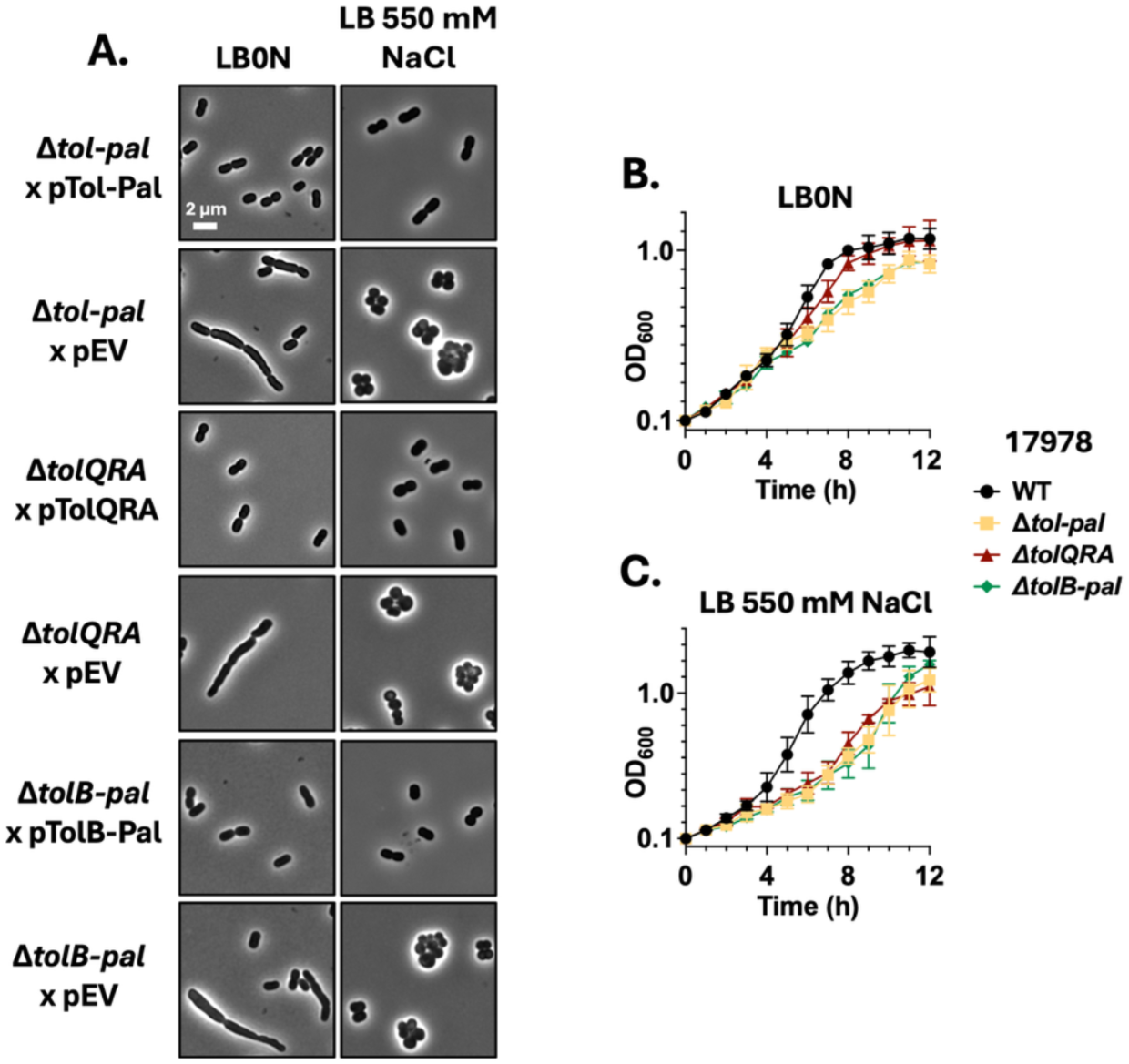
Complementation suppresses osmotic phenotypes in Tol-Pal–deficient cells. Complementation restores envelope organization under osmotic stress. (A) Phase-contrast and HADA labeling of complemented and empty-vector control strains grown in LB0N or LB supplemented with 550 mM NaCl. (B) Growth curves of WT and Tol-Pal–deficient strains under reduced osmolarity. (C) Growth curves under hyperosmotic conditions.

**Figure S5.**
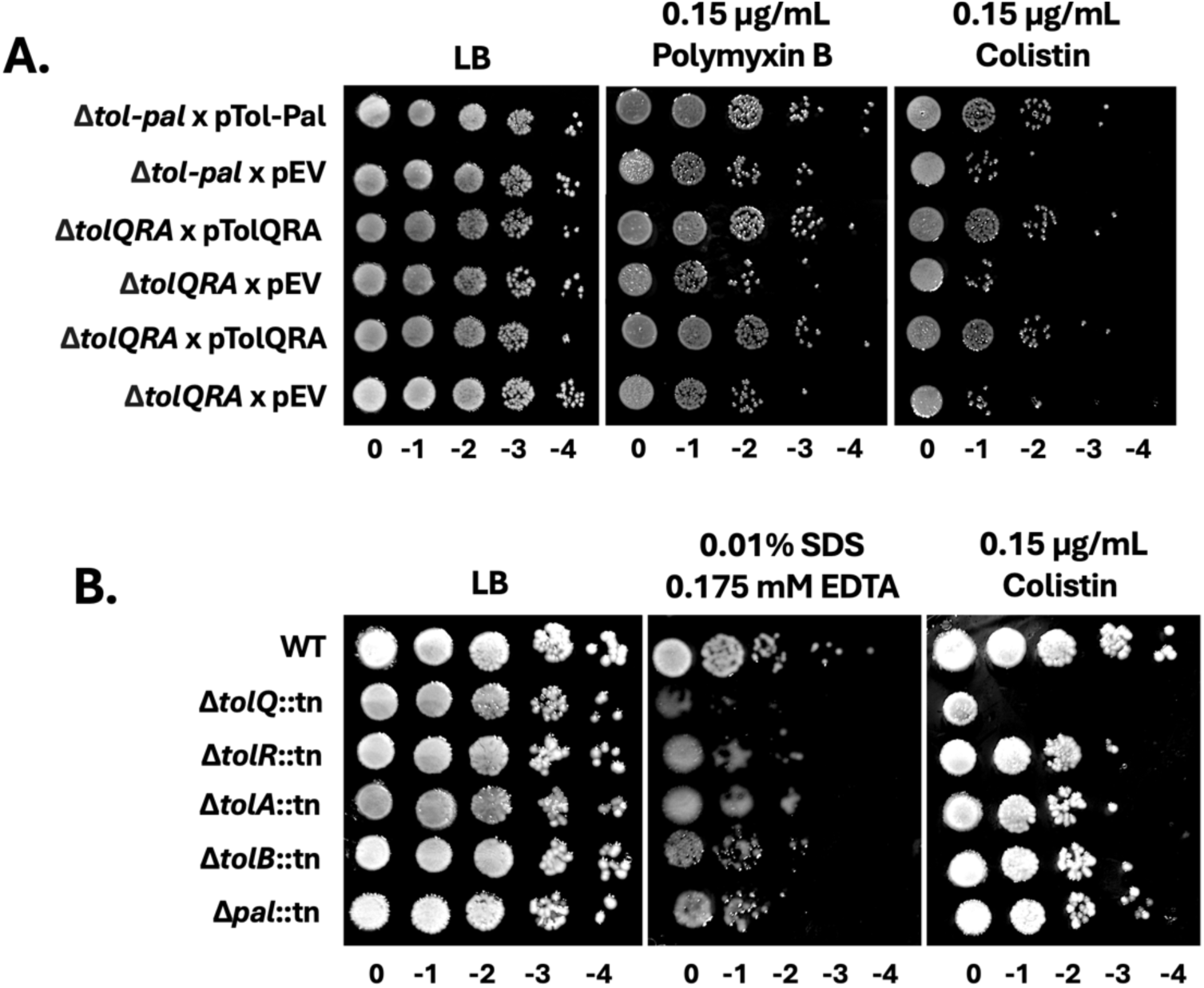
OM barrier defects are conserved across lineages and rescued by complementation. Sensitivity assays in AB5075 and complemented strains. (A) Spot-dilution assays of complemented ATCC 17978 Δ*tol–pal* and Δ*tolQRA* strains compared with empty-vector controls. (B) Spot-dilution assays of AB5075 transposon mutants assessing sensitivity to SDS–EDTA, and colistin.

**Figure S6.**
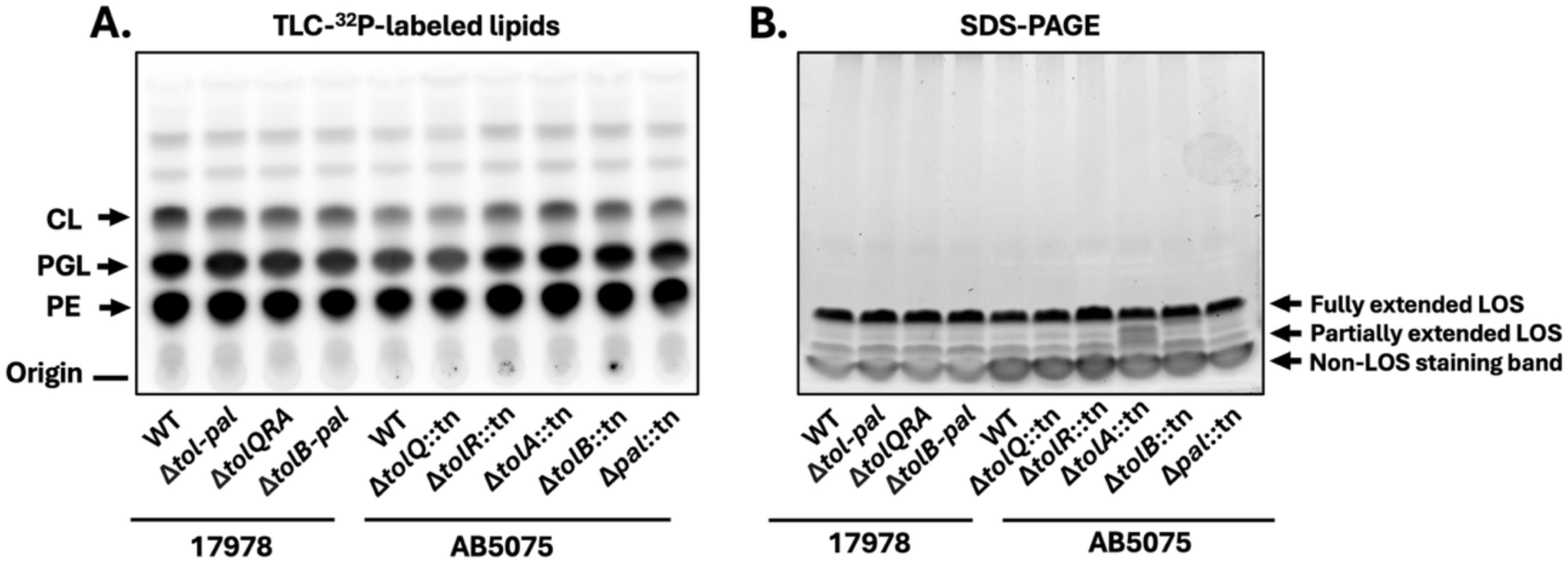
Global phospholipid and LOS profiles are unchanged in Tol-Pal mutants. Envelope lipid composition in Tol-Pal–deficient strains. (A) Thin-layer chromatography analysis of ³²P-labeled phospholipids from WT and Tol-Pal–deficient strains in ATCC 17978 and AB5075 backgrounds. PE, phosphatidylethanolamine; PGL, phosphatidylglycerol; CL, cardiolipin. (B) SDS-PAGE analysis of LOS profiles from WT and Tol-Pal–deficient strains. Images shown are representative of three independent experiments. Quantification of phospholipid species is presented in the Supplementary Table 1 and total LOS levels is presented in Supplementary Table 2.

**Figure S7.**
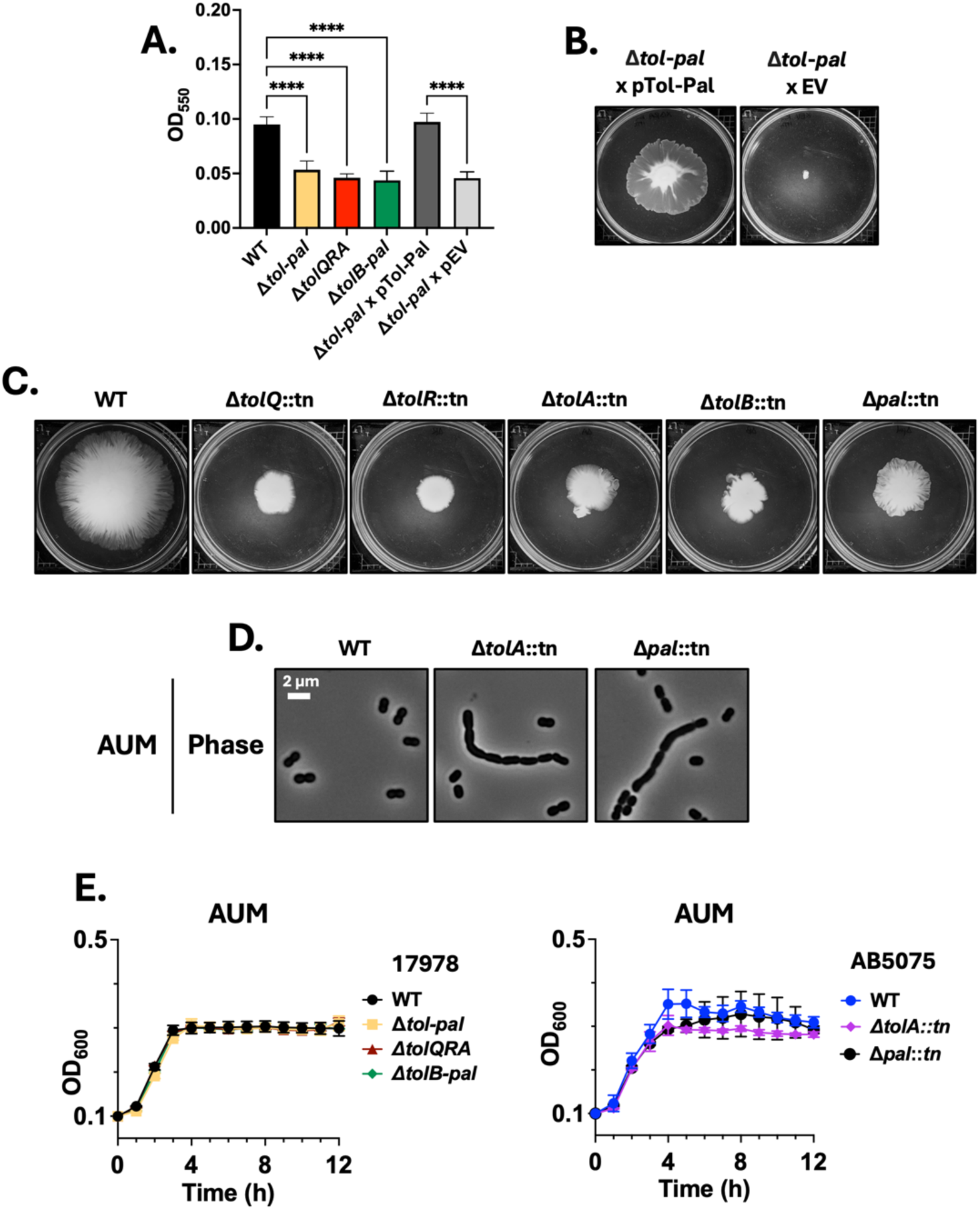
Tol-Pal contributes to fitness and morphology in host-like environments. Tol-Pal-dependent phenotypes in environmental and host-associated assays. (A–C) Biofilm formation and motility assays for complemented and empty-vector control strains. (D) Phase-contrast imaging of AB5075 Tol-Pal transposon mutants grown in AUM. (E) Growth curves of WT and Tol-Pal–deficient strains in AUM.

## Supplemental Figure legends

**Table S1.** Quantification of membrane phospholipid composition in wild-type and Tol-Pal mutant strains.

**ATCC 17978**
| **Lipid** | **WT** | **$\Delta tol-pal$** | **$\Delta tolQRA$** | **$\Delta tolB-pal$** |
| --- | --- | --- | --- | --- |
| **CL** | 23.75 ± 1.2 | 21.51 ± 1.4 | 21.35 ± 1.3 | 21.49 ± 1.5 |
| **PGL** | 35.15 ± 0.9 | 35.31 ± 1.0 | 34.49 ± 0.9 | 34.09 ± 1.1 |
| **PE** | 41.10 ± 1.4 | 43.18 ± 1.6 | 44.16 ± 1.5 | 44.42 ± 1.7 |

**AB5075**
| **Lipid** | **WT** | **$\Delta tolQ$** | **$\Delta tolR$** | **$\Delta tolA$** | **$\Delta tolB$** | **$\Delta pal$** |
| --- | --- | --- | --- | --- | --- | --- |
| **CL** | 18.72 ± 1.1 | 17.68 ± 1.3 | 19.34 ± 1.2 | 20.01 ± 1.1 | 19.88 ± 1.3 | 19.15 ± 1.2 |
| **PGL** | 34.17 ± 0.9 | 34.53 ± 1.0 | 36.47 ± 1.2 | 37.03 ± 1.1 | 36.36 ± 1.0 | 36.33 ± 1.1 |
| **PE** | 47.11 ± 1.5 | 47.79 ± 1.4 | 44.19 ± 1.6 | 42.96 ± 1.5 | 43.76 ± 1.4 | 44.52 ± 1.5 |

**Table S2.** Quantification of LOS abundance in wild-type and Tol-Pal mutant strains.

| **ATCC 17978** |  | **AB5075** |  |
| --- | --- | --- | --- |
| **Strain** | **LOS (mean ± SD)** | **Strain** | **LOS (mean ± SD)** |
| WT | 1.00 ± 0.00 | WT | 1.00 ± 0.00 |
| $\Delta tol-pal$ | 1.07 ± 0.19 | $\Delta tolQ$ | 0.99 ± 0.13 |
| $\Delta tolQRA$ | 1.09 ± 0.14 | $\Delta tolR$ | 1.13 ± 0.34 |
| $\Delta tolB-pal$ | 1.07 ± 0.14 | $\Delta tolA$ | 1.02 ± 0.26 |
| | | $\Delta tolB$ | 1.08 ± 0.10 |
| | | $\Delta pal$ | 1.13 ± 0.26 |

