## Supplemental Material for "Environmental context reveals a conditional role of the Tol-Pal system in envelope organization in *Acinetobacter baumannii*"

### Supplementary Figures

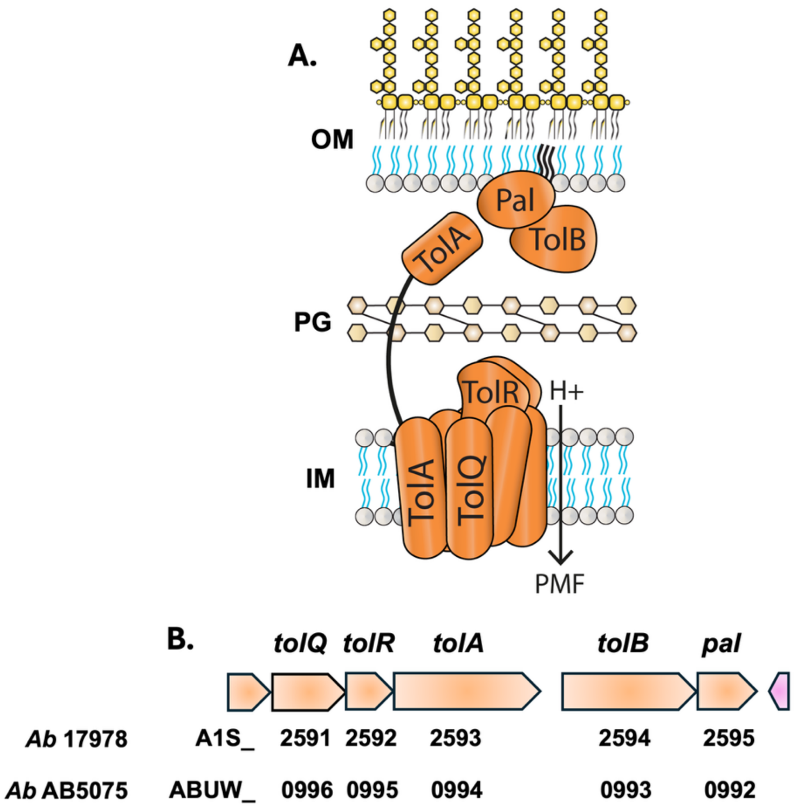

**Figure S1. Organization of the Tol-Pal system in *A. baumannii*.** (A) Schematic representation of the Tol-Pal trans-envelope complex showing the inner membrane (IM) motor components TolQ, TolR, and TolA, which harness the proton motive force (PMF), the periplasmic protein TolB, and the outer membrane (OM) lipoprotein Pal, which associates with peptidoglycan (PG). (B) The corresponding genetic organization of the *tol–pal* locus is, with gene annotations indicated below each locus. The locus is organized into two transcriptional units, *tolQRA* and *tolB–pal*, with gene annotations indicated below each locus.

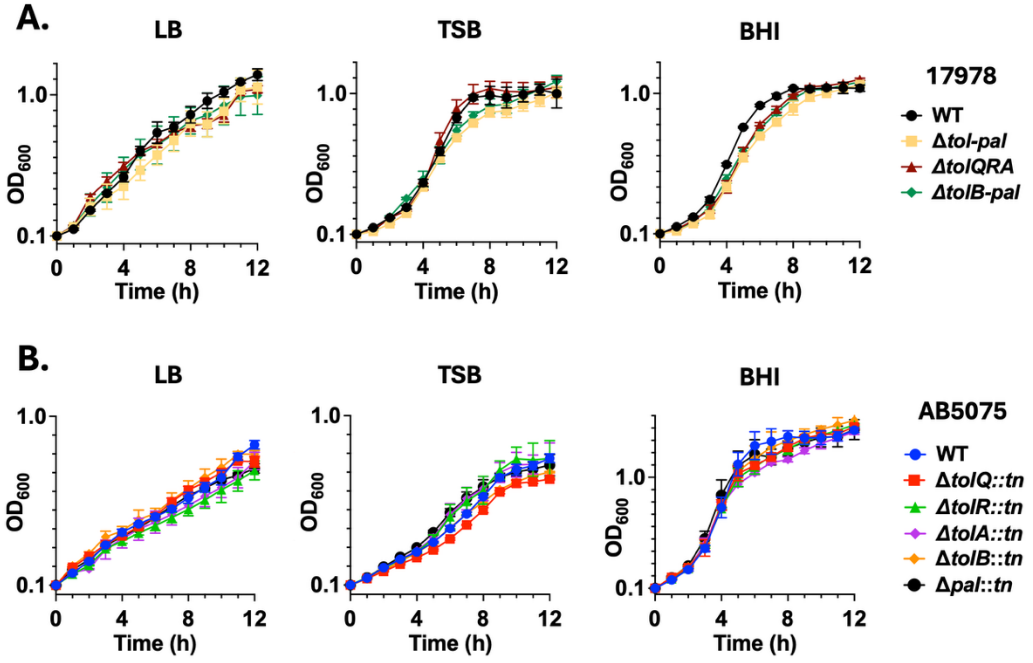

**Figure S2. Tol-Pal deficiency permits sustained population growth under standard laboratory conditions.** (A) Growth of wild-type (WT), Δ*tol–pal*, Δ*tolQRA*, and Δ*tolB–pal* generated in strain ATCC 17978 strains grown in LB, TSB, and BHI. (B) Growth of strain AB5075 WT and transposon mutants disrupting individual Tol-Pal components (*tolQ*, *tolR*, *tolA*, *tolB*, or *pal*) in the same media. Optical density was monitored over time. Error bars represent the standard deviation from three independent biological replicates.

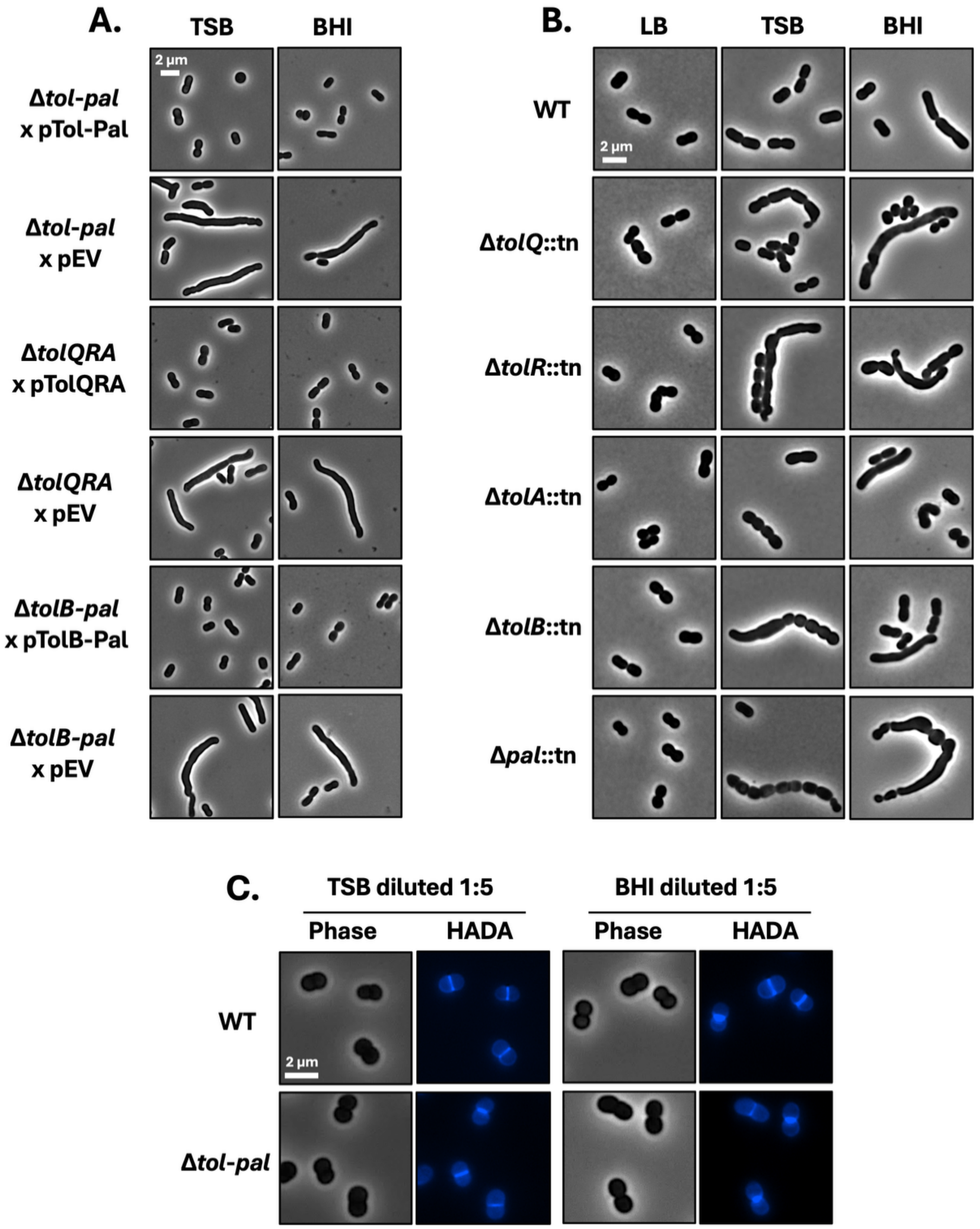

**Figure S3. Genetic complementation restores normal cell morphology, and environmental modulation suppresses Tol-Pal–associated division defects. Role of Tol-Pal is conserved across genetic backgrounds.** (A) Phase-contrast microscopy of *A. baumannii* ATCC 17978 *tol-pal* mutants and complemented strains grown in TSB or BHI. Cell length distributions are shown for complemented Δ*tol–pal*, Δ*tolQRA*, and Δ*tolB–pal* strains expressing the corresponding genes, compared with empty vector controls. Complementation restores wild-type cell morphology and reduces filamentation. (B) Phase-contrast microscopy of AB5075 Tol-Pal transposon mutants grown in LB, TSB, and BHI, showing conserved morphology defects across strain backgrounds and growth conditions. (C) Environmental modulation suppresses Tol-Pal–associated division defects. Phase-contrast and HADA labeling of WT and Δ*tol–pal* cells grown in TSB or BHI diluted 1:5 with water. Medium dilution suppresses filamentation and restores focused mid-cell PG incorporation in Δ*tol–pal* cells. Scale bars, 2 μm.

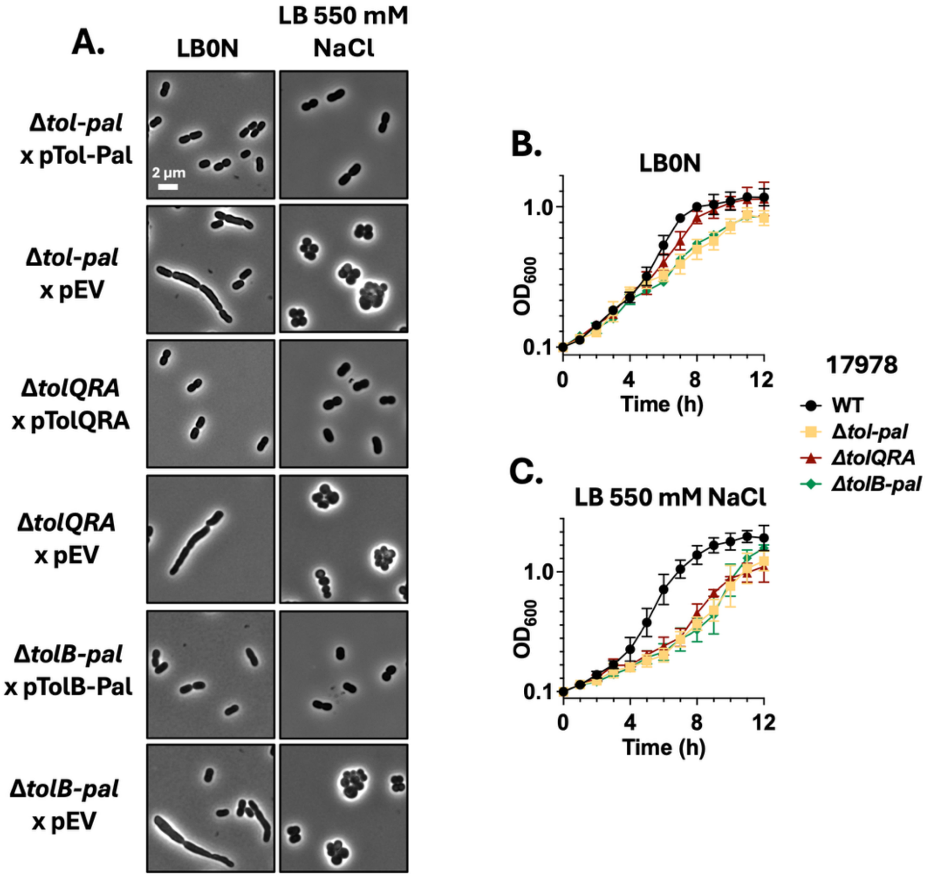

**Figure S4. Complementation suppresses osmotic phenotypes in Tol-Pal–deficient cells.** Complementation restores envelope organization under osmotic stress. (A) Phase-contrast and HADA labeling of complemented and empty-vector control strains grown in LB0N or LB supplemented with 550 mM NaCl. (B) Growth curves of WT and Tol-Pal–deficient strains under reduced osmolarity. (C) Growth curves under hyperosmotic conditions.

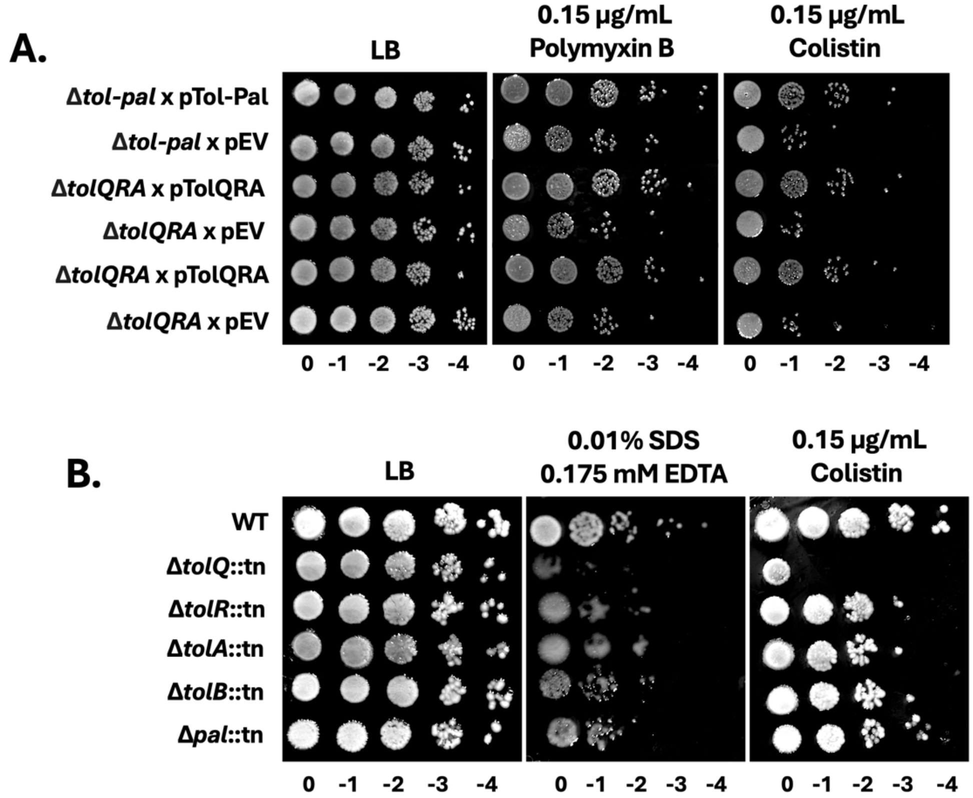

**Figure S5. OM barrier defects are conserved across lineages and rescued by complementation.** Sensitivity assays in AB5075 and complemented strains. (A) Spot-dilution assays of complemented ATCC 17978 Δ*tol–pal* and Δ*tolQRA* strains compared with empty-vector controls. (B) Spot-dilution assays of AB5075 transposon mutants assessing sensitivity to SDS–EDTA, and colistin.

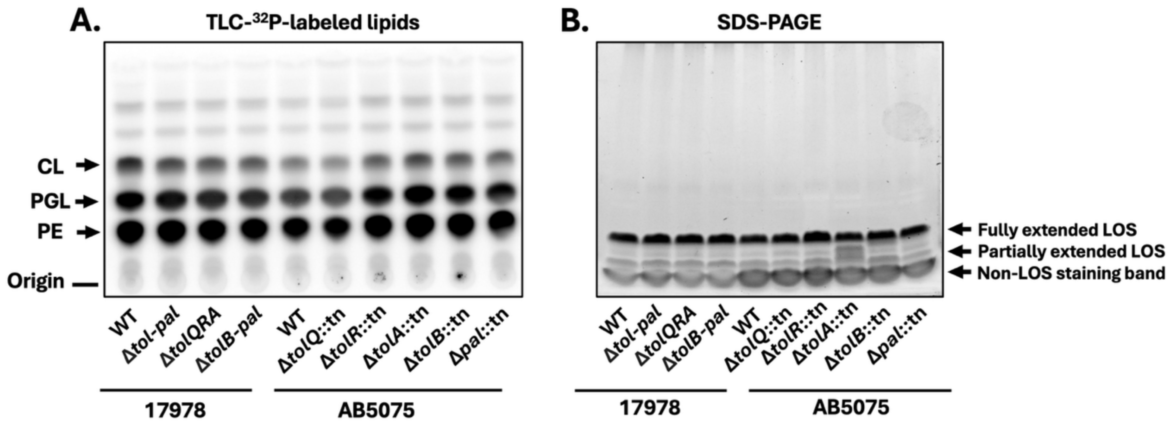

**Figure S6. Global phospholipid and LOS profiles are unchanged in Tol-Pal mutants.** Envelope lipid composition in Tol-Pal–deficient strains. (A) Thin-layer chromatography analysis of $^{32}$P-labeled phospholipids from WT and Tol-Pal–deficient strains in ATCC 17978 and AB5075 backgrounds. PE, phosphatidylethanolamine; PGL, phosphatidylglycerol; CL, cardiolipin. (B) SDS-PAGE analysis of LOS profiles from WT and Tol-Pal–deficient strains. Images shown are representative of three independent experiments. Quantification of phospholipid species is presented in the Supplementary Table 1 and total LOS levels is presented in Supplementary Table 2.

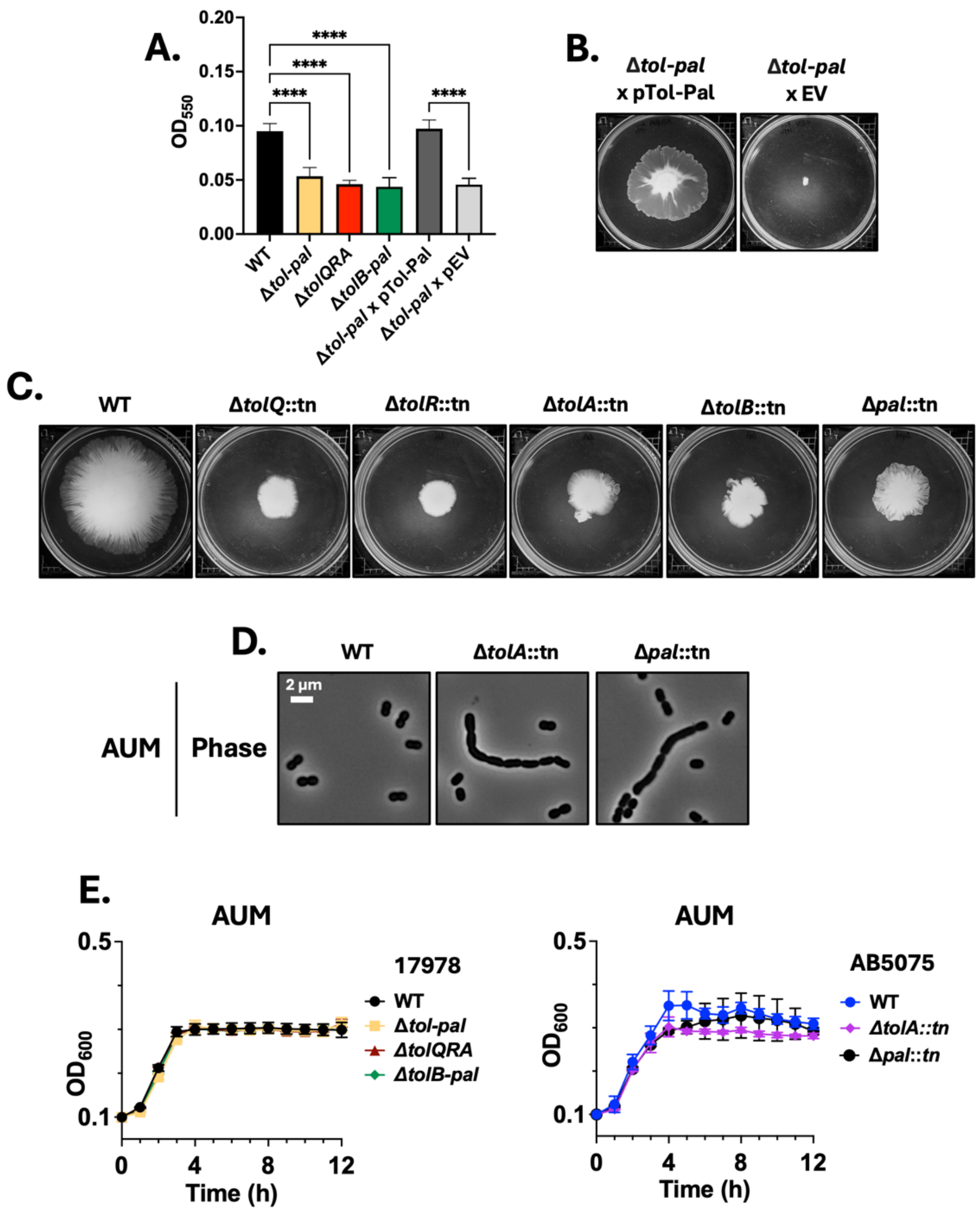

**Figure S7. Tol-Pal contributes to fitness and morphology in host-like environments.** Tol-Pal-dependent phenotypes in environmental and host-associated assays. (A–C) Biofilm formation and motility assays for complemented and empty-vector control strains. (D) Phase-contrast imaging of AB5075 Tol-Pal transposon mutants grown in AUM. (E) Growth curves of WT and Tol-Pal–deficient strains in AUM.

**Supplementary Tables**

**Table S1. Quantification of membrane phospholipid composition in wild-type and Tol-Pal mutant strains.**

**ATCC 17978**

| Lipid | WT | Δ*tol-pal* | Δ*tolQRA* | Δ*tolB-pal* |
|---|---|---|---|---|
| **CL** | 23.75 ± 1.2 | 21.51 ± 1.4 | 21.35 ± 1.3 | 21.49 ± 1.5 |
| **PGL** | 35.15 ± 0.9 | 35.31 ± 1.0 | 34.49 ± 0.9 | 34.09 ± 1.1 |
| **PE** | 41.10 ± 1.4 | 43.18 ± 1.6 | 44.16 ± 1.5 | 44.42 ± 1.7 |

**AB5075**

| Lipid | WT | Δ*tolQ* | Δ*tolR* | Δ*tolA* | Δ*tolB* | Δ*pal* |
|---|---|---|---|---|---|---|
| **CL** | 18.72 ± 1.1 | 17.68 ± 1.3 | 19.34 ± 1.2 | 20.01 ± 1.1 | 19.88 ± 1.3 | 19.15 ± 1.2 |
| **PGL** | 34.17 ± 0.9 | 34.53 ± 1.0 | 36.47 ± 1.2 | 37.03 ± 1.1 | 36.36 ± 1.0 | 36.33 ± 1.1 |
| **PE** | 47.11 ± 1.5 | 47.79 ± 1.4 | 44.19 ± 1.6 | 42.96 ± 1.5 | 43.76 ± 1.4 | 44.52 ± 1.5 |

**Table S2. Quantification of LOS abundance in wild-type and Tol-Pal mutant strains.**

| ATCC 17978 | | AB5075 | |
|---|---|---|---|
| **Strain** | **LOS (mean ± SD)** | **Strain** | **LOS (mean ± SD)** |
| WT | 1.00 ± 0.00 | WT | 1.00 ± 0.00 |
| Δ*tol-pal* | 1.07 ± 0.19 | Δ*tolQ* | 0.99 ± 0.13 |
| Δ*tolQRA* | 1.09 ± 0.14 | Δ*tolR* | 1.13 ± 0.34 |
| Δ*tolB-pal* | 1.07 ± 0.14 | Δ*tolA* | 1.02 ± 0.26 |
| | | Δ*tolB* | 1.08 ± 0.10 |
| | | Δ*pal* | 1.13 ± 0.26 |

**Table S3: Strains and plasmids used in this study.**

| Strain or Plasmid | Description | Reference or Source |
|---|---|---|
| **Strains** | | |
| *A. baumannii* ATCC 17978 | Wild type | (1) |

| | | |
|---|---|---|
| *A. baumannii* ATCC 17978 | *Δtol-pal* | This study |
| *A. baumannii* ATCC 17978 | *ΔtolQRA* | This study |
| *A. baumannii* ATCC 17978 | *ΔtolB-pal* | This study |
| *A. baumannii* ATCC AB5075 | *ΔtolQ::tn26* | (2) |
| *A. baumannii* ATCC AB5075 | *ΔtolR::tn26* | (2) |
| *A. baumannii* ATCC AB5075 | *ΔtolA::tn26* | (2) |
| *A. baumannii* ATCC AB5075 | *ΔtolB::tn26* | (2) |
| *A. baumannii* ATCC AB5075 | *Δpal::tn26* | (2) |
| *E. coli* DH5α | recA1, ϕ80 lacZΔM15, host for cloning | (3) |
| **Plasmids** | | |
| pAT03 | pMMB67EH with FLP recombinase, $Tet^R$ | (4) |
| pAT04 | pMMB67EH with $REC_{Ab}$ system, $Tet^R$ | (4) |
| pKD4 | $Km^R$ | (5) |
| pMMB67EH-Km | pMMB67EH with the $Km^R$ gene from pKD4 inserted into the PvuI site, $Km^R$ | (6) |
| pMMB67EH-Km | pMMB67EH-Km-*tol-pal* (ITPG) | This study |
| pMMB67EH-Km | pMMB67EH-Km-*tol-QRA* (ITPG) | This study |
| pMMB67EH-Km | pMMB67EH-Km-*tolB-pal* (ITPG) | This study |

**Table S4: Primers used in this study.**

| **Oligo Name** | **Sequence (5' to 3')** |
|---|---|
| **Deletion Primers** | |
| 17978 *tol-pal* Kan FRT 5' | ATGGCAACAAACATTGAATCAACCCTGCATATTTCTGACCTTATTTTACAAGCAAGTCCAGTCGTCCAGTTGGTCATGCTGATTT |

| | |
|---|---|
| | TATTGTTAGCATCAATTTTTAGTTGGTACCTGATTGCCAAAGC GATTGTGTAGGCTGGAGCTGCTTCG |
| 17978 *tol-pal* Kan FRT 3' | TTATTTTAATAGAGGAGGAACCGCTTCATAGTTAATTTCAACG CGGCGGTTTTCTTTCCAAGCTGATTCATCATGGCCAGGATTA ACAGGCGCTTCTTTACCATAACTTACAGCTTCAAGTTGCTATA TCCTCCTTAGTTCCTATTCCG |
| 17978 *tol-pal* 5' confirm | CGCTATATGGAACGTACCCGTACAG |
| 17978 *tol-pal* 3' confirm | GACTTGAGCCAACTCAGGTGTAAG |
| 17978 *tolQRA* Kan FRT 5' | ATGGCAACAAACATTGAATCAACCCTGCATATTTCTGACCTTA TTTTACAAGCAAGTCCAGTCGTCCAGTTGGTCATGCTGATTT TATTGTTAGCATCAATTTTTAGTTGGTACCTGATTGCCAAAGC GATTGTGTAGGCTGGAGCTGCTTCG |
| 17978 *tolQRA* Kan FRT 3' | TTATTGTGCTCTGAAAGTAGAGGTAACGCTTCGCGCTTCTCG TCTTGCATCAGGGTCCGATGGCATTGGGTAGGGAGCAGACG CCTGAATTGCAGCTTTGATACTTGCATCCAATGCATCATCTCA TATCCTCCTTAGTTCCTATTCCG |
| 17978 *tolQRA* 5' confirm | CGCTATATGGAACGTACCCGTACAG |
| 17978 *tolQRA* 3' confirm | CTTGCCAATCAGATGCTTGAATCTGG |
| 17978 *tolB-pal* Kan FRT 5' | ATGAAAACGAGCCGCAAACACCTTCTAGCCTTAACGCTACTT ACAGCATTAAGTCCTCTTGCACCAACAGCTGCCTTTGCTCAA TTACATTTAGAGATTGCAAAAGCACCTGATCAGGCGCCTAAA GCGATTGTGTAGGCTGGAGCTGCTTCG |
| 17978 *tolB-pal* Kan FRT 3' | TTATTTTAATAGAGGAGGAACCGCTTCATAGTTAATTTCAACG CGGCGGTTTTCTTTCCAAGCTGATTCATCATGGCCAGGATTA ACAGGCGCTTCTTTACCATAACTTACAGCTTCAAGTTGCTATA TCCTCCTTAGTTCCTATTCCG |
| 17978 *tolB-pal* 5' confirm | GGATGTTCCTACAGGAAGCAGTGG |
| 17978 *tolB-pal* 3' confirm | GACTTGAGCCAACTCAGGTGTAAG |
| **Complementation Primers** | |
| 17978 *tol-pal* BamHI 5' | CGC<u>GGATCC</u>ATGGCAACAAACATTGAATCAACCCTG |
| 17978 *tol-pal* SalI 3' | CGC<u>GTCGAC</u>TTATTTTAATAGAGGAGGAACCGCTTCATAG |
| 17978 *tolQRA* BamHI 5' | CGC<u>GGATCC</u>ATGGCAACAAACATTGAATCAACCCTG |
| 17978 *tolQRA* SalI 3' | CGC<u>GTCGAC</u>TTATTGTGCTCTGAAAGTAGAGGTAACGC |
| 17978 *tolB-pal* BamHI 5' | CGC<u>GGATCC</u>ATGAAAACGAGCCGCAAACACC |
| 17978 *tolB-pal* SalI 3' | CGC<u>GTCGAC</u>TTATTTTAATAGAGGAGGAACCGCTTCATAG |
| **Transposon mutant primers** | |

| | |
|---|---|
| AB5075 *tolQ::tn26* 5' | CTTGCTTGCTTCTGGTGAGGTTGAG |
| AB5075 *tolQ::tn26* 3' | GTACCAGCATTACATCGATATAAGGCACG |
| AB5075 *tolR::tn26* 5' | GCGAAAGTGTCTACTCTGATCGTGC |
| AB5075 *tolR::tn26* 3' | GGTTGTTCAGGTGGTTTACTCAAGCC |
| AB5075 *tolA::tn26* 5' | CAAGTTGGCCTTCTGACAGAGCC |
| AB5075 *tolA::tn26* 3' | GACTGGTGATTTTGTAGCATAGTTACATGC |
| AB5075 *tolB::tn26* 5' | GCATGTAACTATGCTACAAAATCACCAGTC |
| AB5075 *tolB::tn26* 3' | GTTGTACCTGTAGTTGCCGTTGTTG |
| AB5075 *pal::tn26* 5' | CCGTATGAATTTACCGAGTGAACAAGG |
| AB5075 *pal::tn26* 3' | GCATGATTTGATTGAGTCAATGCAACATTCAG |
